# Towards community-driven metadata standards for light microscopy: tiered specifications extending the OME model

**DOI:** 10.1101/2021.04.25.441198

**Authors:** Mathias Hammer, Maximiliaan Huisman, Alex Rigano, Ulrike Boehm, James J. Chambers, Nathalie Gaudreault, Alison J. North, Jaime A. Pimentel, Damir Sudar, Peter Bajcsy, Claire M. Brown, Alexander D. Corbett, Orestis Faklaris, Judith Lacoste, Alex Laude, Glyn Nelson, Roland Nitschke, Farzin Farzam, Carlas S. Smith, David Grunwald, Caterina Strambio-De-Castillia

**Affiliations:** RNA Therapeutics Institute, UMass Medical School, Worcester MA 01605, USA; Program in Molecular Medicine, UMass Medical School, Worcester MA 01605, USA; Janelia Research Campus, Howard Hughes Medical Institute, Ashburn, VA 20147, USA; Institute for Applied Life Sciences, University of Massachusetts, Amherst, MA 01003, USA; Allen Institute for Cell Science, Seattle, WA 98109, USA; The Rockefeller University, New York, NY 10065, USA; Instituto de Biotecnologıa, Universidad Nacional Autonoma de Mexico, Cuernavaca, Morelos, 62210, México; Quantitative Imaging Systems LLC, Portland, OR 97209, USA; National Institute of Standards and Technology, Gaithersburg, MD 20899, USA; Advanced BioImaging Facility (ABIF), McGill University, Montreal, Quebec, H3G 0B1, Canada; Department of Physics and Astronomy, University of Exeter, Exeter, EX4 4QL, UK; BCM, Univ. Montpellier, CNRS, INSERM, Montpellier, France; MIA Cellavie Inc., Montreal, Quebec, H1K 4G6, Canada; Bioimaging Unit, Newcastle University, Newcastle upon Tyne, NE2 4HH, UK; Life Imaging Center and BIOSS Centre for Biological Signaling Studies, Albert-Ludwigs-University Freiburg, Freiburg, 79104, Germany; Delft Center for Systems and Control & Department of Imaging Physics, Delft University of Technology, Delft, 2628 CN, The Netherlands

**Keywords:** Imaging, microscopy, metadata, quality control, calibration, standards, data-formats, reproducibility, open microscopy

## Abstract

Digital light microscopy provides powerful tools for quantitatively probing the real-time dynamics of subcellular structures. While the power of modern microscopy techniques is undeniable, rigorous record-keeping and quality control are required to ensure that imaging data may be properly interpreted (*quality*), reproduced (*reproducibility*), and used to extract reliable information and scientific knowledge which can be shared for further analysis (*value*). Keeping notes on microscopy experiments and quality control procedures ought to be straightforward, as the microscope is a machine whose components are defined and the performance measurable. Nevertheless, to this date, no universally adopted community-driven specifications exist that delineate the required information about the microscope hardware and acquisition settings (i.e., microscopy “data provenance” metadata) and the minimally accepted calibration metrics (i.e., microscopy quality control metadata) that should be automatically recorded by both commercial microscope manufacturers and customized microscope developers. In the absence of agreed guidelines, it is inherently difficult for scientists to create comprehensive records of imaging experiments and ensure the quality of resulting image data or for manufacturers to incorporate standardized reporting and performance metrics. To add to the confusion, microscopy experiments vary greatly in aim and complexity, ranging from purely descriptive work to complex, quantitative and even sub-resolution studies that require more detailed reporting and quality control measures.

To solve this problem, the **4D N**ucleome Initiative (4DN) (*1, 2*) Imaging Standards Working Group (IWG), working in conjunction with the **B**io**I**maging **N**orth **A**merica (BINA) Quality Control and Data Management Working Group (QC-DM-WG) (*3*), here propose light Microscopy Metadata specifications that scale with experimental intent and with the complexity of the instrumentation and analytical requirements. They consist of a revision of the Core of the Open Microscopy Environment (OME) Data Model, which forms the basis for the widely adopted Bio-Formats library (*4*–*6*), accompanied by a suite of three extensions, each with three tiers, allowing the classification of imaging experiments into levels of increasing imaging and analytical complexity (*7, 8*). Hence these specifications not only provide an OME-based comprehensive set of metadata elements that should be recorded, but they also specify which subset of the full list should be recorded for a given experimental tier. In order to evaluate the extent of community interest, an extensive outreach effort was conducted to present the proposed metadata specifications to members of several core-facilities and international bioimaging initiatives including the **E**uropean **L**ight **M**icroscopy **I**nitiative (ELMI), **G**lobal **B**io**I**maging (GBI), and **E**uropean **M**olecular **B**iology **L**aboratory (EMBL) - **E**uropean **B**ioinformatics **I**nstitute (EBI). Consequently, close ties were established between our endeavour and the undertakings of the recently established **QUA**lity Assessment and **REP**roducibility for Instruments and Images in **Li**ght **Mi**croscopy global community initiative (*9*). As a result this flexible 4D**N**-**B**INA-**O**ME (NBO namespace) framework (*7, 8*) represents a turning point towards achieving community-driven Microscopy Metadata standards that will increase data fidelity, improve repeatability and reproducibility, ease future analysis and facilitate the verifiable comparison of different datasets, experimental setups, and assays, and it demonstrates the method for future extensions. Such universally accepted microscopy standards would serve a similar purpose as the Encode guidelines successfully adopted by the genomic community (*10, 11*). The intention of this proposal is therefore to encourage participation, critiques and contributions from the entire imaging community and all stakeholders, including research and imaging scientists, facility personnel, instrument manufacturers, software developers, standards organizations, scientific publishers, and funders.

## 2 - INTRODUCTION

The reproducibility crisis affecting the biological sciences is well-documented (*12*–*16*). In the field of light microscopy, it can only be addressed if all published images are accompanied by complete descriptions of experimental procedures, biological samples, microscope hardware specifications, image acquisition settings, image analysis parameters and metrics detailing instrument performance and calibration (*4, 9, 13, 17*–*20*). This complete description, also known as Image Metadata, consists of any and all information about an imaging experiment that ensures its rigorous interpretation, reproducibility and re-use, and should be recorded alongside the actual image data in the file header or in supplemental files (*21*). A fully developed metadata model would provide for consistent tracking of crucial information pertaining to the quality, reproducibility and scientific value of image data, and will allow the communication and comparison of such information in a Findable, Accessible, Interoperable, and Reproducible (FAIR) manner (see also Text Box 1 in: Huisman et al., 2021) (*21, 22*). However, as microscopy has evolved from a tool that generates purely descriptive or illustrative data to primary quantitative data acquired with ever more sophisticated and complex instruments, our practices to record this quantitative data and metadata faithfully and reproducibly have not kept up. Moreover, while the Open Microscopy Environment (OME) consortium (*5, 6, 23*–*25*) has made significant advances with the development of the OME Data Model which, together with the widely-used Bio-Formats library (*4*), serves as the only available *de facto* specification for accessing and exchanging image data, the field of light microscopy still lacks universally accepted standards for imaging data and specifications for metadata. The want of a consensus has resulted in an out-of-control growth of proprietary and/or incompatible image file formats and metadata capture practices.

This manuscript is intended to launch a community-driven way forward to break the impasse. Specifically, it puts forth scalable specifications for light Microscopy Metadata developed jointly by the **4D N**ucleome Initiative (4DN) (*1, 2*) **I**maging Standards **W**orking **G**roup (IWG) and by the **B**io**I**maging **N**orth **A**merica (BINA) **Q**uality **C**ontrol and **D**ata **M**anagement **W**orking **G**roup (QC-DM-WG) (*3, 7, 8*). In order to foster widespread adoption of these 4D**N**-**B**INA-**O**ME (NBO namespace) (*26*) specifications framework (Figure 1A, magenta bubble), parallel work is being conducted to develop user-friendly and when possible automated metadata collection tools (*13, 27*–*30*); shared metadata storage best practices (Figure 1A, yellow bubble) (*31*–*34*); as well as sustainable specifications to switch from proprietary formats for image data into common, cloud-ready OME Next-Generation File Formats (NGFF, Figure 1A, blue bubble) (*35, 36*). Importantly, all of these activities are carried out in the context of the newly launched **QUA**lity Assessment and **REP**roducibility for Instrument and Images in **Li**ght **Mi**croscopy (QUAREP-LiMi) initiative (quarep.org; (*9, 37*) and involve several key members of the community, including microscope users, custodians, and manufacturers, imaging scientists, national and global bioimaging organizations, bio-image informaticians, standards organizations, and scientific publishers. The light Microscopy Metadata guidelines proposed here articulate along three orthogonal axes (Figure 1B):

**Figure 1:**
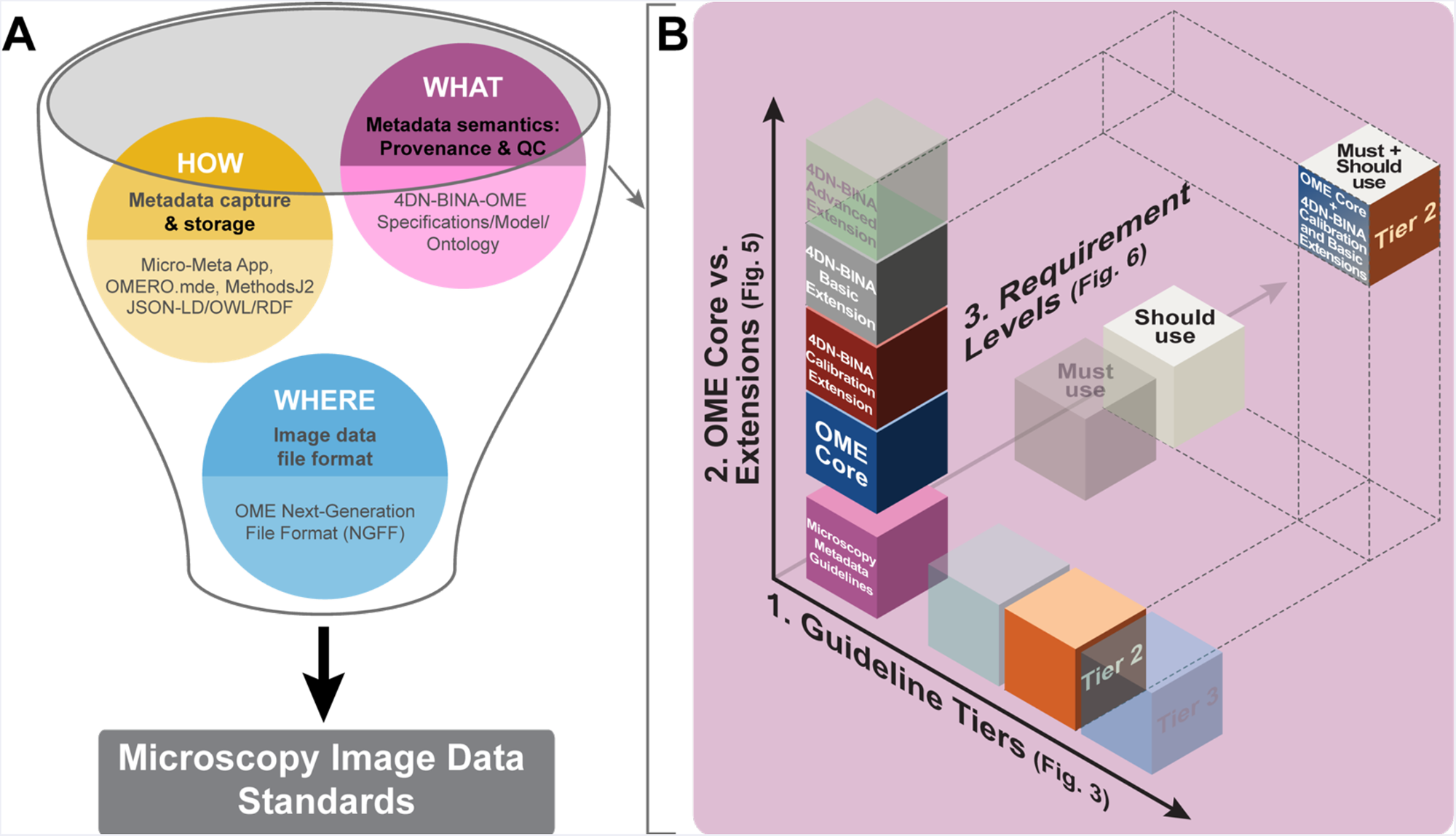
The definition of community-driven Microscopy Image Data Standards requires three complementary components and needs a flexible framework to manage complexity. **A)** The establishment of community-driven Microscopy Image Data Standards implies parallel development on three interrelated fronts: 1) Community-driven specifications for what ‘data provenance’ information and quality control metrics are essential for rigor, reproducibility, and reuse and should therefore be captured in Microscopy Metadata (*magenta bubble*). 2) Shared rules for how the (ideally) automated capture, representation and storage of Microscopy Metadata should be implemented in practice (*yellow bubble*). Last but not least, 3) Next-Generation File Formats (NGFF) where the ever-increasing scale and complexity of image data and metadata would be contained for exchange(35,36); *blue bubble*). All these provisions must be compatible with the FAIR principles of data stewardship(22), maximize consistency, flexibility, extensibility, and adapt to ever-growing experimental and technical complexity. **B)** Guidelines for what hardware specifications, image acquisition settings, and quality control metrics should be captured in microscopy metadata to ensure rigor, reproducibility, and reuse articulate along three complexity axes: 1) Guideline Tiers: The use of one of the three guidelines Tiers that scale with reporting requirements of experiments of increasing complexity (Figure 3 and Table II). 2) OME Core vs. Extensions: The use of the Core of the OME Data Model vs. one or more of the 4DN-BINA extensions. 3) Requirement Level: The use of Required (*Must use*) vs. Recommended (*Should use*) metadata fields. In the example shown, the selected combination in which all available metadata fields of the OME Core + 4DN-BINA Calibration and Basic extensions used at Tier 2 would be appropriate to describe an experiment in which a wide-field microscope is used to capture a time series that was acquired to capture the dynamics viral particles trafficking within infected cells.

**Table I:**
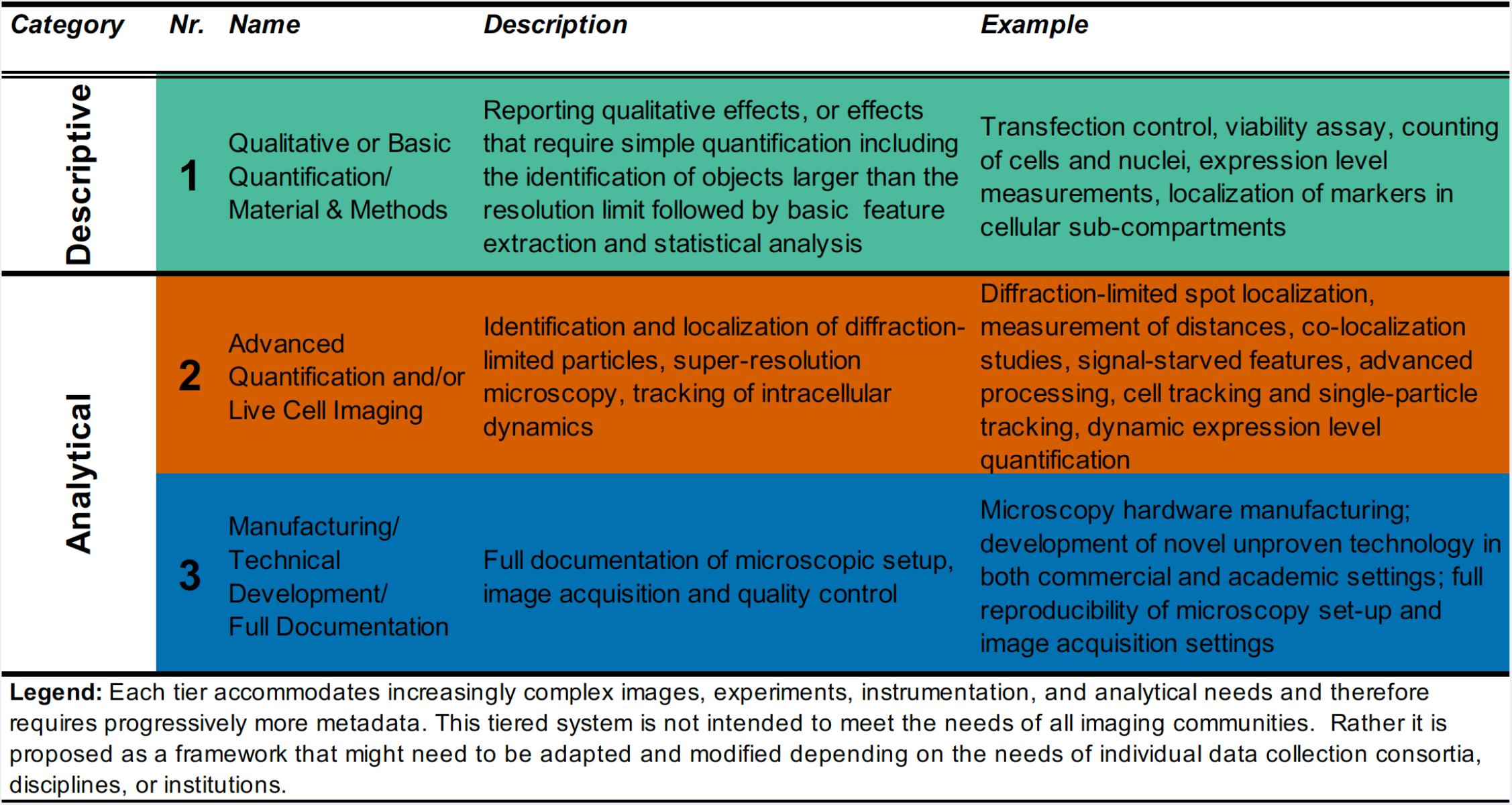
Tiers for light microscopy metadata reporting as proposed by the Imaging Standards Working Group of the 4D Nucleome initiative and by the Quality Control and Data Management Working Group of Bioimaging North America.

**Table II:**
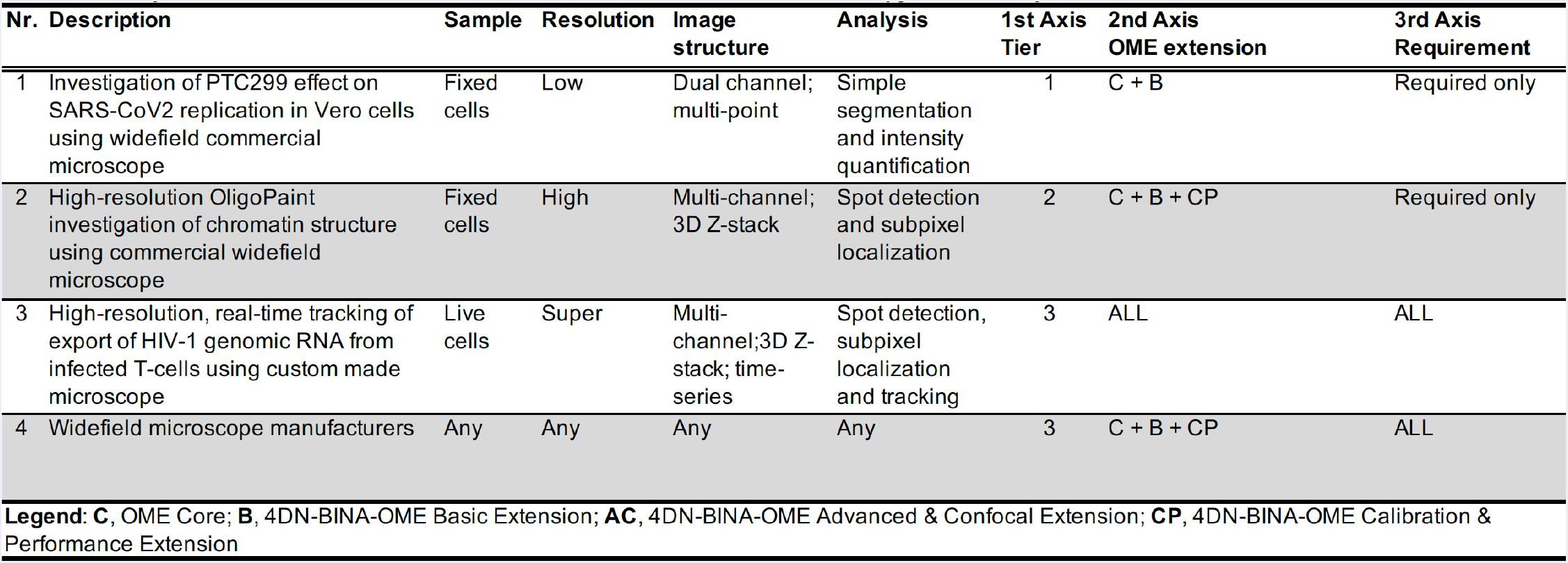
Example utilization of the three axes matrix of the 4DN-BINA -OME Microscopy Metadata Specifications

|  |  |  |  |  |  |  |  |  |
| --- | --- | --- | --- | --- | --- | --- | --- | --- |
| Nr. | Description | Sample | Resolution | Image structure | Analysis | 1st Axis Tier | 2nd Axis OME extension | 3rd Axis Requirement |
| 1 | Investigation of PTC299 effect on SARS-CoV2 replication in Vero cells using widefield commercial microscope | Fixed cells | Low | Dual channel; multi-point | Simple segmentation and intensity quantification | 1 | C + B | Required only |
| 2 | High-resolution OligoPaint investigation of chromatin structure using commercial widefield microscope | Fixed cells | High | Multi-channel; 3D Z-stack | Spot detection and subpixel localization | 2 | C + B + CP | Required only |
| 3 | High-resolution, real-time tracking of export of HIV-1 genomic RNA from infected T-cells using custom made microscope | Live cells | Super | Multi-channel; 3D Z-stack; time-series | Spot detection, subpixel localization and tracking | 3 | ALL | ALL |
| 4 | Widefield microscope manufacturers | Any | Any | Any | Any | 3 | C + B + CP | ALL |

**Legend:** **C**, OME Core; **B**, 4DN-BINA-OME Basic Extension; **AC**, 4DN-BINA-OME Advanced & Confocal Extension; **CP**, 4DN-BINA-OME Calibration & Performance Extension

1. <u>Guideline Tiers</u> - Metadata specification (*7, 8*): A system of adaptable Tiers that spells-out which specific subset of metadata information should be included depending on experimental intent, technical complexity, and image analysis needs.
2. <u>Core model and Extensions</u> - Metadata extension (*26*): An suite of extensions that expand the core of the OME Data Model (*4, 5*) to better capture state-of-the-art transmitted light and widefield fluorescence microscopy (Basic Extension), and confocal and advanced fluorescence modalities (Advanced and Confocal Extension). Importantly to improve the management of quality control, a novel data model for capturing instrument calibration procedures (Calibration and Performance Extension) was developed in close collaboration with QUAREP-LiMi (*9, 37*).
3. <u>Requirement Levels - Metadata inclusion</u>: Inherent flexibility in the inclusion of metadata is built in the model so that specific pieces of information will be considered as “required” (essential for rigor and reproducibility), and “recommended” (to improve image quality and to maximize scientific and sharing value).

The framework of this model is inherently adaptable while providing microscope users and custodians, instrument makers and microscope vendors with a clear and enforceable community-driven mandate for the necessary information to ensure scientific rigor, experimental reproducibility and maximal scientific value.

### 2.1 The metadata challenge in microscopy imaging: the great variability of data formats and metadata reporting practices

The capture and analysis of microscopy images closely depend on the technique utilized to record and measure light. The introduction of photography, fluorescence, computers and digital light detectors drastically improved the objectivity of observations made through light microscopy and changed light microscopy in three profound ways. First, it has allowed the increasingly accurate recording of progressively lower amounts of light, enabling the visualization and quantitative measurement of sub-cellular and single-molecule (SM) events and molecular interactions with high specificity and temporal resolution. Second, the advent of digital image formation and processing has enabled new imaging modalities, such as Confocal Laser Scanning Microscopy (CLSM), and super-resolution (SR) imaging techniques that allow high-resolution imaging of live samples in three dimensions. Third, digital imaging has led to signal processing and computational methods that enable the extraction of quantitative information from images. Despite these advances, the emergence of light (and in particular fluorescence) microscopy as a key quantitation tool for biomedical research, and the employment of ever more sophisticated and complex instruments, practices to record this quantitative data and metadata faithfully and reproducibly have not kept up.

When performing imaging experiments, scientific rigor is inextricably tied to image quality, the reproducibility of experimental results and the extent to which image datasets have a scientific value. Not only can a high-value dataset be used to answer the scientific questions it was intended to address, but it can also be shared, merged with other datasets, integrated with other data types and further analyzed to answer new questions. Deriving valuable information from images is completely dependent on the consistent recording and storage of “data provenance” information that capture sample preparation and labeling, microscope hardware specification and image acquisition details, on the quantitative assessment of the optical, excitation, detection, and mechanical properties (including error estimation) of the microscope, and on an intimate knowledge of the analysis procedures used to extract quantitative information from the images. Hence the proliferation of quantitative light microscopy techniques has not only opened new scientific landscapes but has also exacerbated the existing challenges of quality control and reproducibility.

A typical light microscopy experiment includes three (sometimes integrated) major steps (Figure 2): 1) Sample Preparation, i.e., all preparative steps resulting in the samples to be imaged; 2) Image Acquisition, i.e., image formation and recording; and 3) Image Analysis, i.e., the post-acquisition processing and quantification of images. Each procedure within these steps can add considerable variability to the final data. Thus, to document all possible sources of uncertainty, microscopy images need to be accompanied by Image Metadata (*21*) describing all information about the imaging experiment that allows the evaluation, interpretation, reproducibility, and comparison of the actual image data (i.e., quantitative values associated with the image pixels; Figure 2, Pixel Image Data). This will include comprehensive information about all aspects of the microscopy experiment from experimental treatment and sample preparation (Figure 2A, Experimental and Sample Metadata) to microscope hardware specifications, image acquisition settings, image structure, and instrument performance metrics (Figure 2A, Microscopy Metadata), as well as details about any image analysis procedures that were subsequently employed (Figure 2A, Analysis Metadata) (*38*–*40*). Although the OME Data Model, coupled with the ubiquitous Bio-Formats image file format conversion library (*4, 5*), has served as the *de facto* exchange format for image data, it has not as yet evolved into a much needed community mandated Microscopy Image Data Standard (*4, 5*). In addition to support for extensions to capture technological advancements, a mature standard would include: 1) universal image data formats defining the container *<u>where</u>* data is stored (*35*); 2) specifications of *<u>what</u>* information about the imaging experiment should be captured; and 3) standards for *<u>how</u>* the metadata should be captured, managed and stored (Figure 1).

**Figure 2:**
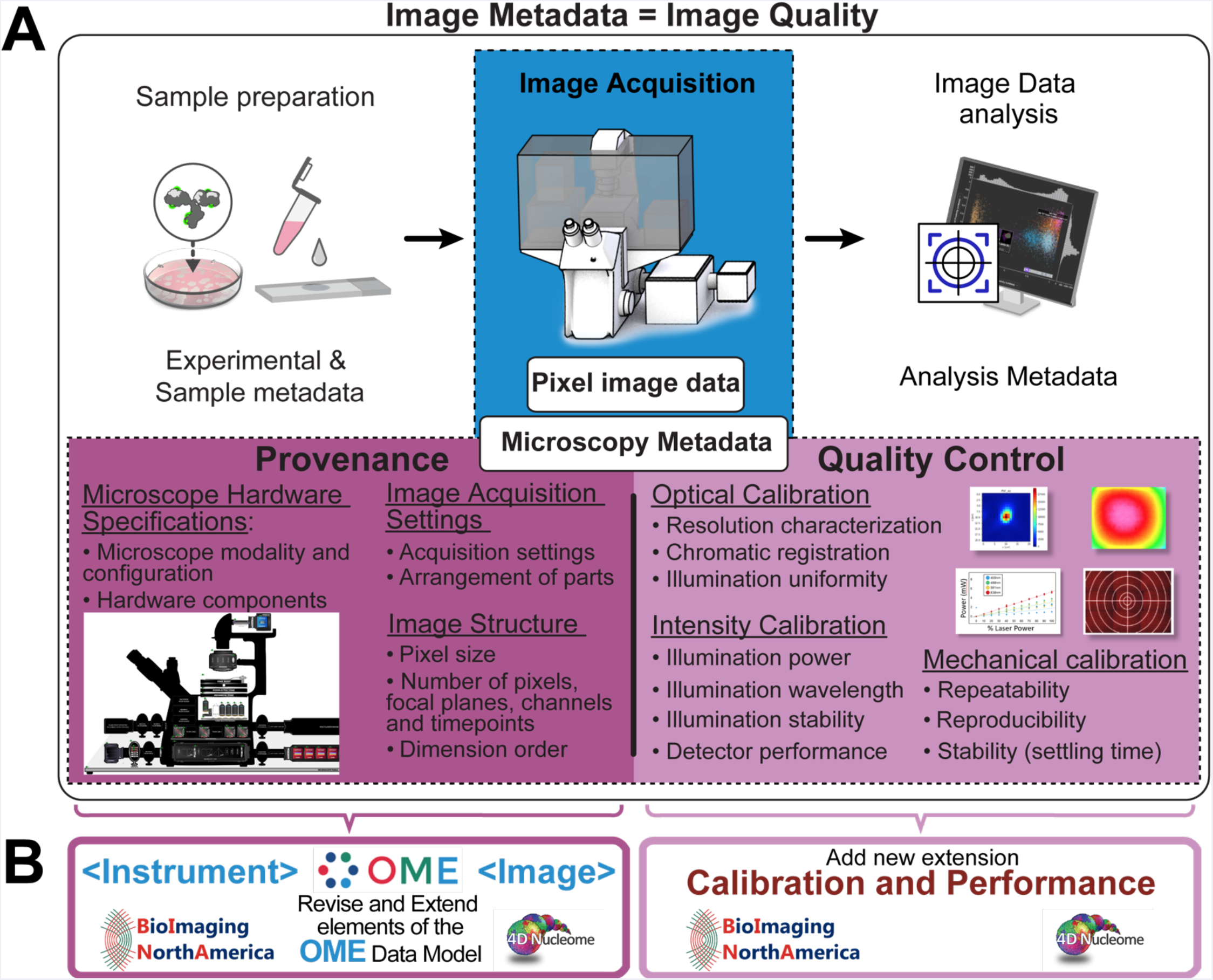
Light Microscopy Metadata is essential for the assessment, interpretation, reproducibility, comparison, and re-use of the results of microscopy experiments. **A)** A typical bio-imaging experiment consists of three main phases at the center of which is the production of image data: 1) Sample preparation, 2) Image Acquisition, and 3) Image Analysis. Each of these phases involves a host of different procedures and techniques and must be properly documented and validated to maximize the value, quality, and reproducibility of the resulting image data. Metadata is data that provides information about other data. **Image Metadata** consists of any and all information that allows the numerical values stored in individual pixels that form image data (blue box, *Pixel Image Data*) to be evaluated, interpreted, reproduced, found, cited, and re-used as established by measurable data quality criteria (i.e., FAIR principles) (*22*). While the importance of Sample Preparation, and Image Analysis metadata cannot be overstated, the discussion here centers specifically around Light **Microscopy Metadata**, defined as the metadata that documents the process of Image Acquisition using a Light Microscope and the quality of the resulting Image Data. When it is properly utilized, carefully maintained, and operated following consistent experimental conditions, a light microscope is expected to display measurable, finite, and repeatable accuracy and precision. It follows that the relative impact of microscopy can be minimized through proper record-keeping and instrument calibration. As such, Microscopy Metadata can be subdivided in two categories: microscopy “provenance” metadata (MPM) includes information about the microscope hardware and the settings that were applied to the instrument during image acquisition; while quality control metadata (QCM) includes metrics that quantitatively assess the performance of the microscope and the quality of image data and are obtained through the execution of specifically designed Optical, Intensity (split in Excitation and Detector) and Mechanical calibration procedures. **B)** In order to capture Microscopy Metadata, the 4DN-BINA light microscopy metadata scalable specifications advanced here were designed to revise and extend the OME Data Model (4,5), which serves as the *de facto* specification for the exchange of image data and metadata. Specifically, Data Provenance metadata capturing Microscope Hardware Specifications and Image Acquisition Settings are stored to revise and extend the `<INSTRUMENT>` and the `<IMAGE>` elements of the OME Data Model, respectively. On the other hand, Quality Control metadata is stored utilizing the specifically designed Calibration and Performance extension of the same model.

Additional challenges in record-keeping for microscopy experiments include the following:

1. **Light microscopy is employed to address a diverse range of complex biological questions**. This has led to the development of a vast array of adaptable microscopy modalities, each requiring different metadata to be reported as well as diverging quality control approaches, which poses a notable challenge when choosing the correct validation method for the experiment at hand.
2. The **working conditions, theoretical performance, and capabilities of the microscope are difficult to assess** and are often unknown by the average user.
3. The **relevant hardware or software metadata can be difficult to retrieve** from available documentation and the user is often unaware of how varying specific parameters might affect the imaging results.
4. The **paucity of automation and intuitive software tools** make record-keeping unduly burdensome, forcing experimental biologists to choose between scientific rigor and productivity.
5. The **lack of universal and enforceable image file format and metadata standards** results in an unacceptable variability in the information provided by microscope manufacturers alongside images. In addition, the need for raw data files to be converted into other formats prior to interpretation and comparison often yields a significant loss of metadata, or, worse still, inadvertently compromises the data during the conversion process.

It is worth noting that among all experimental steps described in Figure 2, the image acquisition step contributes the most manageable and quantifiable stage, as long as the microscope and imaging system are properly documented, maintained, and operated. Consequently, the development of community-sanctioned standards for the compilation of Microscopy Metadata encompassing, Microscopy data Provenance Metadata (MPM), which documents the process of image acquisition and the structure of the resulting image (Figure 2A, *Provenance Metadata*), and Microscopy Quality-control Metadata (MQM), which captures calibration procedures and metrics (Figure 2, *Quality-Control Metadata*) is not only essential for image data quality, reproducibility, and scientific and sharing value, but should be readily attainable (*21*).

### 2.2 The importance and potential pitfalls of standardization

The value of Microscopy Image Data Standards (Figure 1) has been widely recognized, resulting in important efforts to establish best performance testing and instrument calibration practices (*41*–*52*), to unify data-submission requirements from journals (*53*) and to produce the exchange format between image data and metadata that forms the basis for this current work (*4, 5, 54, 55*). Nonetheless, because the existing efforts lack normative value it remains challenging to determine which parameters are relevant to a given technique and imaging experiment and best practice recommendations are often ignored due to their perception as too expensive, complicated and cumbersome.

Much would thus be gained from harmonizing the reporting standards in light microscopy. First, it would facilitate the documentation of any microscopy-based protocol, minimize error, and quantify residual uncertainty associated with each step of the procedure (Figure 2). This, in turn, would provide a wealth of valuable contextual information - collectively referred to as “data provenance” (*56, 57*) – that would greatly increase the scientific and sharing value of the data. Such details would enable the reliable evaluation of scientific claims based on imaging data, facilitate comparisons within and between experiments, allow for reproducibility, and maximize the likelihood that data can be collated and analyzed by other scientists using current and future image processing and analysis methods. Furthermore, the increasing availability of public image repositories (e.g., Movincell (*58*), Image Data Resource - IDR (*55*), Electron Microscopy Public Image Archive - EMPIAR (*59*), Bioimage Archive (*60*), Allen Cell Explorer (*61*), the Cochin Image Database (*62*), the Cell Image Library (*63*), the RIKEN Systems Science of Biological Dynamics database - SSBD (*64*), and the NIH CELL Image Library (*65*)), will undoubtedly increase the need for community-wide documentation and quality control standards, which can adapt to new technologies. As a first step in this direction (*20*) the Recommended Metadata for Biological Images (REMBI) (*66*) guidelines were recently proposed that would maximize the possibility of making bioimaging datasets available to other researchers in a timely manner, consistent with the FAIR principles (Findable, Accessible, Interoperable and Reusable) (*22*), and thus amenable for reuse.

Despite offering innumerable advantages, standardization also has its pitfalls. First, it can massively increase the administrative burden associated with imaging experiments. Second, a lack of flexibility can severely limit the type of data that can be stored. Since it is impossible to know *a priori* the complexity and diversity inherent to experimental details and imaging modalities that are yet to be developed, it is critical that any proposed set of guidelines that are designed to serve as a basis for a sustainable community standard are defined utilizing technology that meets strict extensibility requirements. Because of its inherent extensibility and the solid plans for modernization (see Text Box 1), the OME Data Model (*4, 5*) provides a robust foundation for Microscopy Metadata (Figure 2B) that can be extended by introducing information that is not yet covered (including experimental specific metadata, modality specific metadata, quality control metadata and analysis-specific metadata). As these extensions (*39, 67, 68*) become more commonly used, they too can then be incorporated into the core using community announcements and related vetting processes to meet expanding community needs.

## 3 - A THREE-DIMENSIONAL MATRIX OF 4DN-BINA-OME MICROSCOPY METADATA GUIDELINES

Since a one-size-fits-all solution for Microscopy Metadata requirements is clearly not tenable, we propose a framework in which microscopy documentation and quality control requirements are organized along three axes that are largely orthogonal to each other (Figure 1B). The first axis is based on the observation that different types of experiments have different reporting and quality control requirements based on technical complexity, experimental design, and image analysis needs. Hence, requirements along this axis are subdivided into Tiers depending on the three criteria listed above (Figure 1B, Guideline Tiers; Figure 3, Table I and Supplemental Table I). The second axis starts with and extends the OME Data Model (*4, 5*) with additional metadata components that are introduced based on the microscopic modality (e.g., epifluorescence vs. confocal microscopy) and accommodates expansion as new technologies are developed that are covered neither by the core nor by the currently proposed extensions (Figure 1B, OME Core vs. Extensions; Figure 5). Last but not least, the third axis grades documentation requirements based on whether each piece of information is essential for rigor and reproducibility (Must use), or recommended to improve image quality and for maximizing scientific and sharing value (Should use; Figure 1B, Requirement Levels; Figure 6). The existence of these three axes will allow institutions, funding agencies, consortia and scientific publishers to define best practices for light microscopy experiment documentation while concomitantly allowing individual scientists to find an appropriate position on the guideline matrix that both matches their needs and remains compatible with community-mandated guidelines. As an example, Table II lists where representative experiments would fall within the Microscopy Metadata guideline matrix (Figure 1B) (*7, 8*)..

**Figure 3:**
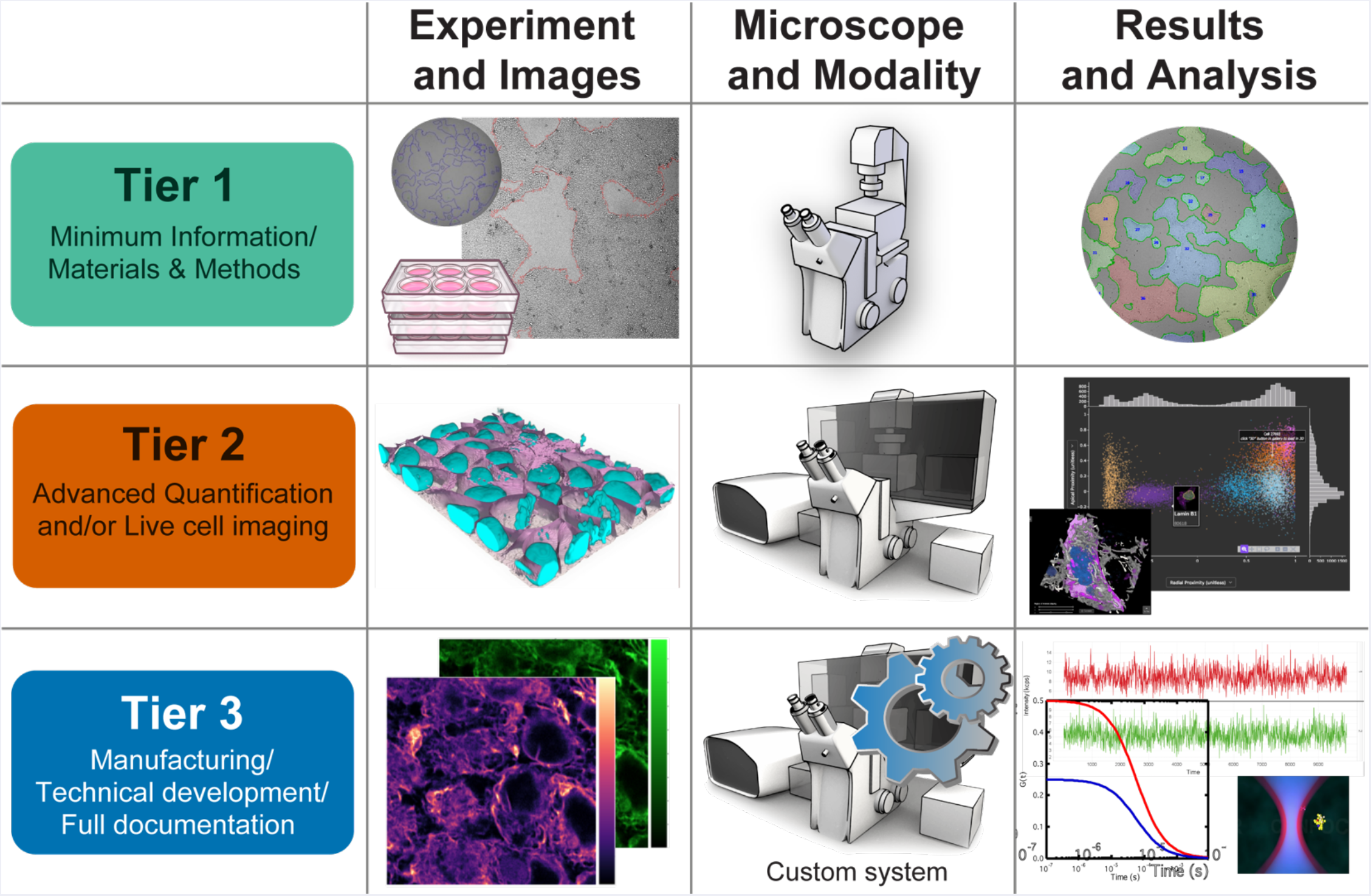
Scaling light microscopy metadata guidelines with experimental, technical, and analytical intent and complexity minimize recordkeeping burden while maximizing value, quality, and reproducibility of image data. Given the complexity of experimental design, technical requirements and analysis procedures, a one-size-fits-all solution for microscopy metadata requirements is unreasonable. In this perspective, a graded system for metadata requirements would minimize the burden of collecting metadata for each experiment while maximizing the opportunities for reproducibility, evaluation, analysis, and comparison. We, therefore, propose a flexible system in which different imaging communities (i.e., consortia, individual research institutions, individual fields of knowledge) would define sets of *a priori* criteria whereby microscope hardware and imaging experiments are classified based on <u>Experiment and Image</u> complexity, <u>Microscope</u> technology and imaging <u>Modality</u>, and <u>Results and Analysis</u> requirements. Specifically, the Tiers levels presented here reflect the recommendations of the 4DN and BINA WGs and should be intended as an initial example of how different imaging experiment types could be placed on a complexity scale to facilitate the collection of the most appropriate minimum set of metadata required for reproducibility and comparison of each category.

**Figure 4:**
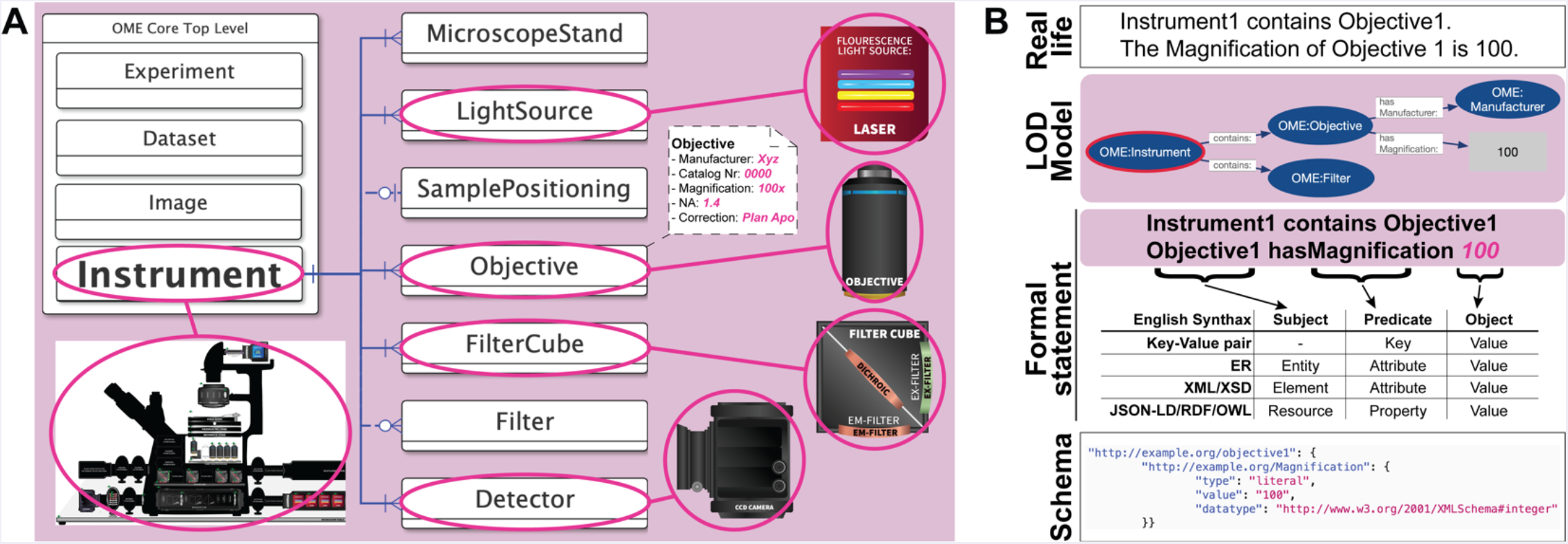
A data model is a schematic representation of reality that can be utilized to organize data. A) A useful way to represent metadata that captures not only the individual attribute and their values but also the often-complex relationships between different real-world components is to build abstract models for the data that account for the components (i.e., Entities) of the system, the attributes that need to be recorded for each component, and the relationship between components. The Entity-Relationship (ER) diagram (70) is a well-known formalism for developing an appropriate data model. In the depicted example an ER schema is used to represent the hardware components of a microscope Instrument. In this representation, nodes (white solid lines boxes) represent individual Entities such as for example Light Source, Objective, Filter Cube and Detector; edges (blue lines) represent Relationships between components (i.e., an Instrument has a Light Source), and field associated with each node represent the data or metadata the model aims to collect about each component (i.e., Objective Manufacturer, Magnification, Numerical Aperture etc.). Attributes are listed as key-value pairs within a dashed-line box and for clarity they are only listed for Objective. Relationships may be required (*solid lines*) or optional (*dashed lines*) and might have different cardinalities: 1 to 1; 1 to 1 … ∞; 1 to 0 … 1; 1 to 0 … ∞. In order to interpret the schema, one typically starts from the `<INSTRUMENT>`, follows one of the blue lines to a connected Entity and states: “A microscope Instrument has one or more Light Sources, and employs one or more Objectives including one built by Nikon, that has 100x Magnification and 1.4 Numerical Aperture”. B) ER diagrams are adequate to provide a bird-eye view of the entire system. However, for a model to serve as the basis for the implementation of essential metadata capture and management tools, it is necessary to represent the data model (*Model*) in a formalized schema (*Schema*). Extensible Markup Language (XML) Schema Definition (XSD) is a very common method to represent a model schema in a machine-readable manner. However, XSD is based on a “closed world” paradigm and makes it difficult to extend data models. An alternative method, which is sanctioned by the World Wide Web Consortium (W3C; https://www.w3.org/), makes use of JSON-Linked Data (JSON-LD) to serialize Resource Description Framework (RDF) triples conforming to Web Ontology Language (OWL) ontologies to build extensible Linked Open Data (LOD) graph models. In practice, human readable statements describing the data (*Real life*) are first rendered into a graphical representation (i.e., LOD graph; *Model)* and then parsed to provide more formalized statements (*Formal statement*). Finally, they are coded using formal schema languages (*Schema*). Depicted is a JSON-LD serialization of RDF/OWL.

**Figure 5:**
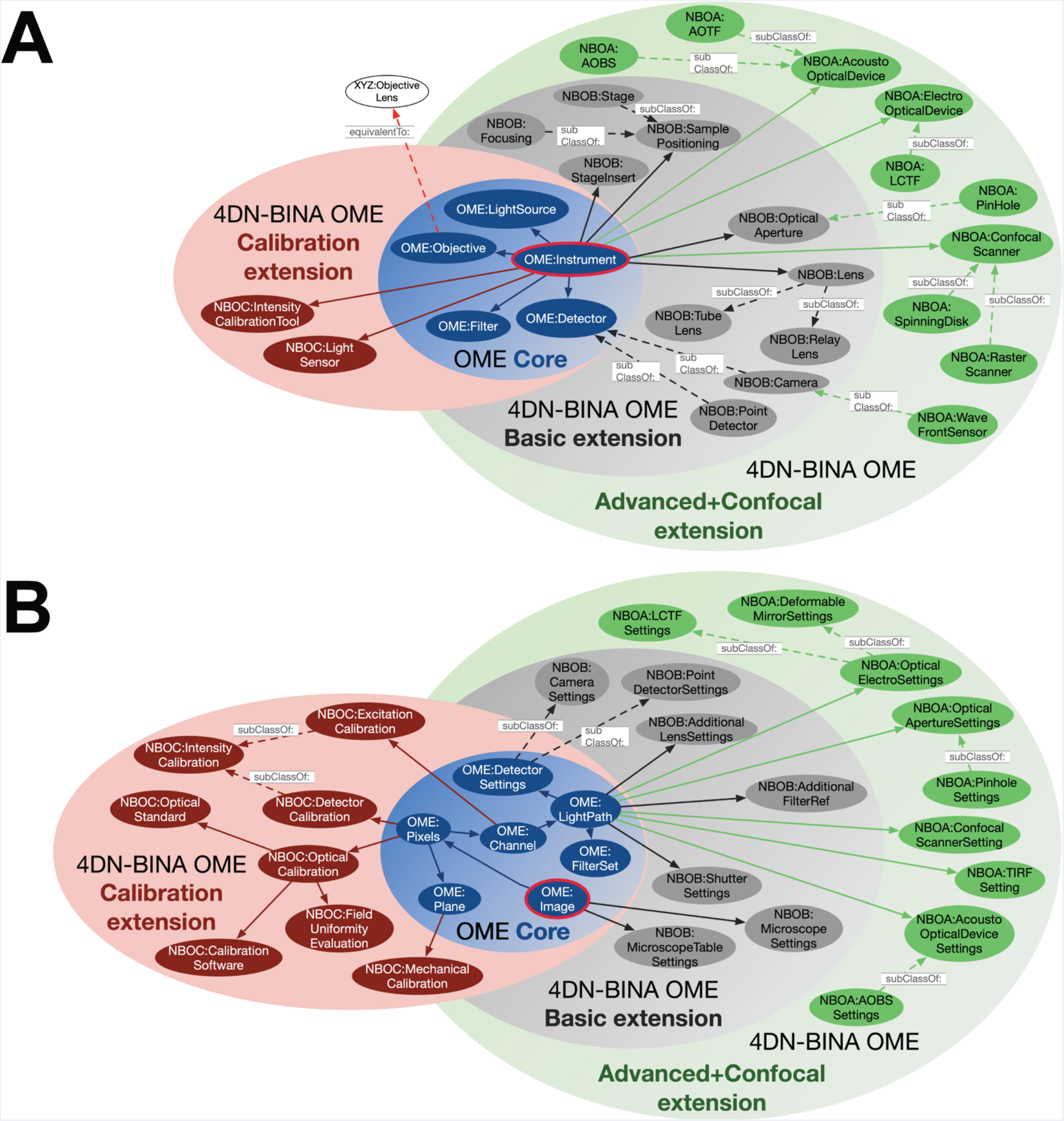
The 4DN-BINA Light Microscopy Metadata Guidelines extends the core of the OME Data Model. A) The OWL/RDF semantic modelling technology stack developed for the World Wide Web allows the construction of flexible data models using the Linked Open Data and “core/extension” paradigms. The 4DN-BINA community-driven extensions of the OME Data Model collectively belong to a new namespace called **4d<u>N</u>-<u>B</u>ina-<u>O</u>me (NBO)**. In turn, novel NBO model elements belong to three different extension packages, which can be used or not depending on the needs of specific imaging communities. A) Portrayed is a Venn diagram depicting the relationship of the `<INSTRUMENT>` element and its sub-elements (*OME: Instrument, red bordered blue circle*) of the Core of the OME Data Model (*blue set*), with sub-elements present in the <u>Calibration and Performance</u> (*Calibration extension, pink set* containing *NBOC maroon circles*), <u>Basic</u> (light *grey set* containing *NBOB dark grey circles*), and <u>Advanced and Confocal</u> (*green set* containing *NBOA green circles*) 4DN-BINA-OME extensions. The schema presented here is not intended to be comprehensive and only includes a small subset of the classes that compose the model. Abbreviations: AOBS, Acousto-Optical Beam Splitter; AOTF, Acousto-Optical Tunable Filter; LCTF, Liquid-Crystal Liquid Filter. B) The OWL/RDF semantic modelling technology stack developed for the World Wide Web allows the construction of flexible data models using the Linked Open Data and “core/extension” paradigms. The 4DN-BINA community-driven extensions of the OME Data Model collectively belong to a new namespace called **4D<u>N</u>-<u>B</u>ina-<u>O</u>me (NBO)**. In turn, novel NBO model elements belong to three different extension packages, which can be used or not depending on the needs of specific imaging communities. B) A) Portrayed is a Venn diagram depicting the relationship of the `<IMAGE>` element and its sub-elements (*OME: Image, red bordered blue circle*) of the Core of the OME Data Model (*blue set*), with sub-elements present in the <u>Calibration and Performance</u> (*Calibration extension, pink set* containing *NBOC maroon circles*), <u>Basic</u> (light *grey set* containing *NBOB dark grey circles*), and <u>Advanced and Confocal</u> (*green set* containing *NBOA green circles*) 4DN-BINA-OME extensions. The schema presented here is not intended to be comprehensive and only includes a small subset of the classes that compose the model.

**Figure 6:**
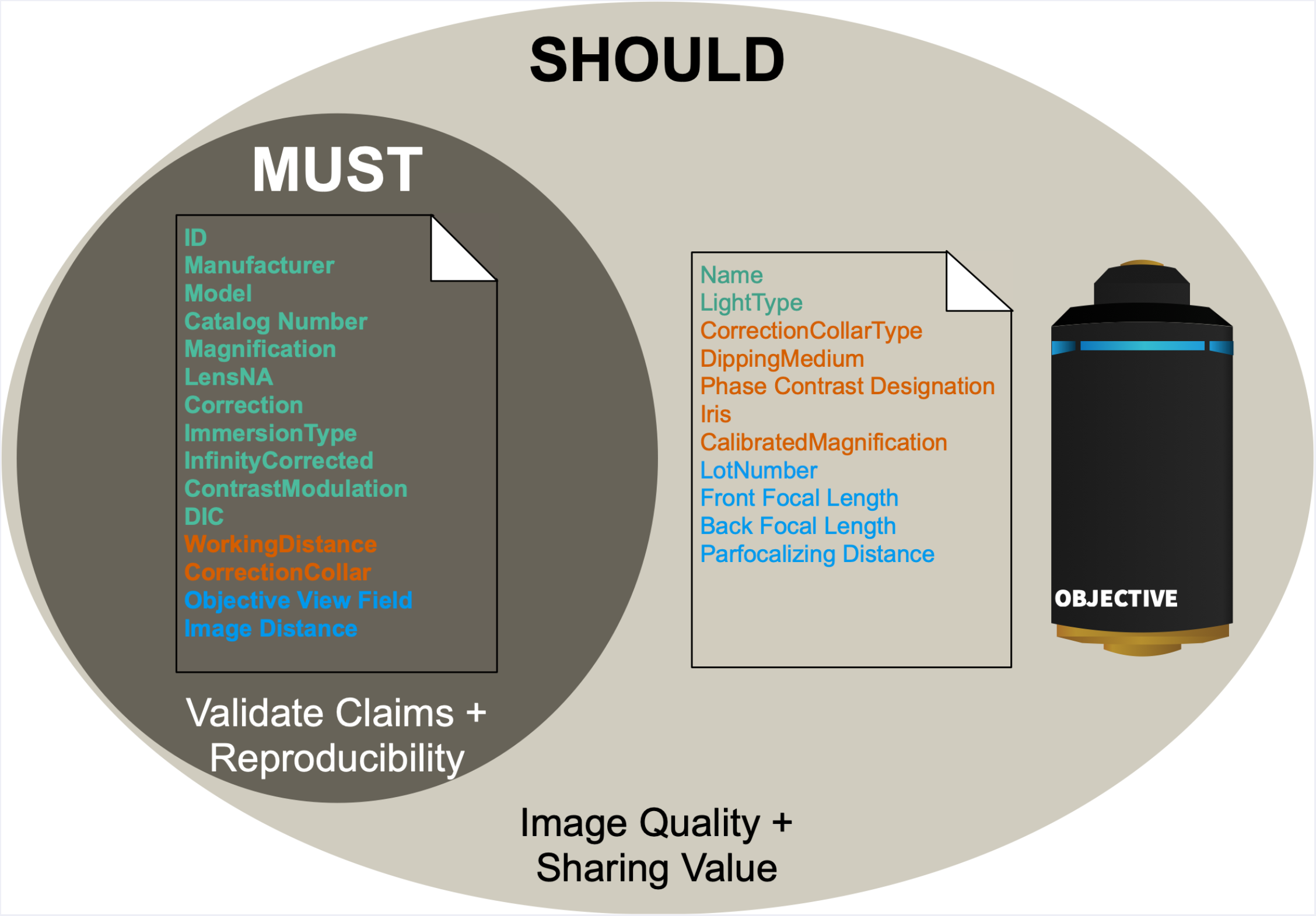
The third axis of the 4DN-BINA-OME light Microscopy Metadata Guidelines adds further flexibility to minimize imaging experiment documentation burden. Depicted is a Venn diagram representing an example referring to the tiered attributes that are required to document the characteristics of an objective lens, and that are stored in the`<OBJECTIVE>` element found in the OME Core and 4DN-BINA Basic extension. These attributes are subdivided into *required* (*MUST*) vs. *recommended* (*SHOULD*) categories to increase documentation flexibility and minimize the administrative and record-keeping burden associated with the documentation of light microscopy experiments. The Tier (Figure 3) each field belongs to is color coded using the same colors utilized throughout the manuscript. Specifically, Tier 1, *Green*. Tier 2, *Orange*. Tier 3, *Blue*.

### 3.1 The first axis: a tier-based system of guidelines for light Microscopy Metadata

To achieve rigor and reproducibility, increasingly elaborate imaging experiments require additional metadata on top of those required for more basic experiments. On this account, a graded system for metadata requirements is not only appropriate, but it also minimizes the burden of collecting metadata for each experiment whilst maximizing the opportunities for rigor, reproducibility, evaluation, analysis, and comparison. We envision a flexible system in which different imaging communities (i.e., individual research institutions, individual fields of knowledge or research consortia) would define their own sets of criteria whereby microscope hardware and imaging experiments are classified in Tiers based on experimental and image complexity, microscope technology and imaging modality, and analytical requirements. Hence the tiered system of guidelines presented here (Figure 3; Table I and Supplemental Table I) (*8*). should be considered as an example of how different imaging experiment types could be placed on a complexity scale to facilitate the collection of the most appropriate minimum set of metadata required for reproducibility and comparison of each category. We expect that this system will naturally evolve organically to incorporate new imaging modalities.

A robust, maximally useful, and efficient metadata standard would be tailored around the different reporting requirements of experiments of increasing complexity. We suggest here a system composed of one Descriptive (Tier 1) and two Analytical (Tiers 2 and 3) tiers (Figure 3; Table I and Supplemental Table I) (*8*), in which imaging instrumentation and datasets are classified based on the following sets of criteria:

1. Are results amenable to visual interpretation or is advanced image analysis (e.g., sub-pixel spot localization) required for the full understanding of results?
2. Are biological samples fixed or alive during acquisition?
3. Are any parts of the quantitative microscopy pipeline (microscope instrument, acquisition modality and image analysis) relying on novel, rather than fully established technology?
4. Is the data provenance and quality control metadata tracked, documented and reported by hardware manufacturers or instrument developers?

Consistent with minimum information principles, the system represents a minimal set of metadata required for each tier, covering only the information relevant for the interpretation of the specific imaging experiment (while more complete information is always allowed and encouraged). As an example, while the proposed standard encompasses information about the sample that directly impacts the imaging conditions (e.g., labeling method, mounting medium), it is not intended to replace the complete description of the experimental procedure from start to end.

#### 3.1.1 Minimum Information required for full documentation → Tier 1

Tier 1 covers experiments that fall into two general sub-categories:

1. <u>Qualitative evaluation</u>: Experiments that require only qualitative assessment of image data for meaningful conclusions to be drawn. Examples include transfection controls, viability tests, or other experiments that serve as minor supporting evidence in a project or manuscript.
2. <u>Simple image processing and analysis on fixed samples</u>: Experiments that are performed on fixed specimens and require simple processing and analyses to support conclusions. This category includes studies that require the identification, counting, intensity and morphometric analyses of features whose size is above the limit of resolution of the system. Examples include cell-counting, the measurement of reporter intensity, the localization of reporters in the nucleus vs. cytoplasm or the estimation of the size and shape of individual cells.

Hence, this descriptive tier does not require metadata describing advanced hardware features of the microscope or quantifying microscope performance. The complete description of metadata fields to be included in Tier 1 is available on GitHub (*7, 8, 26, 69*).

#### 3.1.2 Advanced image analysis, live-imaging and super-resolution -- Tier 2

Advances in microscope technology have been accompanied by an increased dependence on complex image analysis methods. Some imaging techniques require digital image processing and image analysis for the very construction of the images (e.g., assembling individual pixel intensities into the pixel array that constitute an image for CLSM, Structured Illumination Microscopy - SIM, and stochastic SR methods like Photo-Activated Localization Microscopy - PALM, and Stochastic Optical Reconstruction Microscopy - STORM). Others use model-driven data-processing to enhance the resolution of the data (e.g., deconvolution) and improve quantitative accuracy and reliable interpretability by combatting issues such as illumination shading, nonspecific background, limited signal, and optical aberrations. In addition, many imaging experiments require advanced image analysis for extracting quantitative information from the raw data. These techniques increase the usefulness of microscopy data but they often require that the data meets certain criteria for the analysis to be useful and reliable, thus requiring more detailed data provenance and performance calibration information for correct interpretation. Specifically, Tier 2 experiments fall into two general sub-categories:

1. <u>Advanced quantification</u>: Experiments that aim to draw conclusions about features that are near or below the limits of resolution, as well as experiments in which conclusions require post-acquisition reconstitution (i.e., deconvolution). Imaging techniques that use probability-based detection frameworks to function in signal-starved conditions also fall in this category, since they often rely on advanced processing and quantitative analysis. Typical examples include single-molecule (SM) localization microscopy (e.g., SM Fluorescence In Situ Hybridization - FISH), single-particle tracking (SPT) and distance/distribution measurements that aim to achieve SR precision. Because these sophisticated imaging techniques require the optical system to be performing at its theoretical best and often take into account the photophysical behaviors of the fluorophores and the detector, Optical and Intensity Calibration (*21*) are an essential requirement.
2. <u>Live cell imaging</u>: Experiments that use transmitted and fluorescence light microscopy on live specimens whether or not they are destined for advanced processing and analysis. These experiments require detailed information about the environmental conditions, phototoxicity data, and focal and stage stability measurements. Typical examples include applications following the real-time dynamics of cellular events and real-time viral SPT experiments. Because the stability of the system across time is often crucial for reproducibility, Mechanical Calibration (*21*) is also recommended for these experiments.

The capacity of microscope users to provide *bona fide* documentation of microscopy experiments is often limited by the degree to which they are made aware of all parts of the instrument by the manufacturer. Accordingly, Tier 2 represents the most demanding requirement for data obtained by microscope users using well-established microscope instrumentation, processing algorithms and analysis procedures that have been shown to be quantifiable across a range of conditions. The complete description of metadata fields to be included in Tier 2 is available on GitHub (*7, 8, 26, 69*).

#### 3.1.3 Manufacturing, technical development and full documentation → Tier 3

Tier 3 is intended to be used by manufacturers of microscope hardware components and for the developers of pioneering technologies. While it is inherently impossible to design a comprehensive metadata standard for future technologies, all efforts should be made by instrumentation developers and manufacturers to provide any piece of information required, implicitly or explicitly, to interpret a given image formation or data processing method. As such, Tier 3 furnishes manufacturers and developers of new microscope modalities with clear community-sanctioned specifications about what provenance reporting and quality control information should be provided to microscope users to ensure scientific rigor, full reproducibility, and re-use value. The complete description of metadata fields included in Tier 3 is available on GitHub (*7, 8, 26, 69*). Additional method-specific information is expected to be required for most applications.

### 3.2 The second axis: a system of 4DN-BINA-sponsored community-driven OME extensions

In its simplest form, metadata can be easily represented as lists of key-value pairs, where the first term is a descriptive term for a specific attribute and the second term is the value of the attribute, including units for numerical values. However, lists of key-value pairs are often not sufficient to define rich metadata guidelines as they do not allow to capture the often complex relationships between different real-world components and situations to be described. A better method is the development of abstract models for the data that represents the scenario to be described. Ideally, such a data model would account for the components of the system, the attributes that need to be recorded for each component to be fully documented, and the relationship between components (Figure 4A). A useful formalism for developing, describing, and viewing an appropriate data model is the Entity-Relationship (ER) diagram (*70*), which subsequently has to be translated into formalized schemas (Figure 4B) to facilitate implementing metadata capture and management tools.

#### 3.2.1 OME: an informatics framework for biological-image data

Since its launch in 2003 (*54*) the OME initiative has played a leading role in the foundation and development of bio-image informatics (*71*), a new field of biological informatics which concentrates explicitly on building technology for the sharing and dissemination of reproducible and quality controlled biomedical image data. Specifically, the OME Data Model initially published in 2005 (*4, 5*) and its related software implementations (*4, 72*) were developed to provide an informatics framework for the accurate capture, storage and management of rich metadata representations of all the information required for interpreting, comparing, reproducing, sharing and publishing biological microscopy data (Figure 2, *Microscopy Metadata*) independent of a proprietary file format. The OME Data Model is first and foremost implemented in the Bio-Formats library (*4*) which allows the interpretation of the more than 150 different proprietary file formats and their translation into the OME-TIFF and OME-NGFF (*35, 36*) open file formats for biological imaging. This facilitates the transfer of image data between different commercial vendors and open-source software tools thus reducing the barriers between different analytical and data management tools. Finally, the OME Data Model is also implemented in OMERO, a relational database and application server to import, store, process, view and export data (*72*).

One of the OME Data Model’s main purposes is to describe the different hardware components of the microscope, define the light path of each channel and document the settings used for individual image acquisition sessions (i.e., laser power, exposure times and detector gain). Since the physical setup of microscopes tends to be fixed, while imaging settings are typically adjusted for different samples and acquisition sessions, the OME Data Model subdivides Microscopy Metadata into two main sections:

1. The `<Instrument>` **core element** (Figure 2B) describes the imaging instrument and is used to store the relatively static description of a given microscope and its hardware components (e.g., objectives, illumination sources, filters, detectors).
2. The `<Image>` **core element** (Figure 2A) stores the specific instance of an acquired image and, in addition to describing the structure of the, possibly multi-dimensional, image, it documents the image acquisition settings utilized when that image was acquired. To this aim, the `<Image>` data element stores references to specific hardware components defined in the `<Instrument>` alongside any necessary configurations and parameter settings utilized for a given image dataset (e.g., excitation power, filter set and detector gain).

While this structure is very robust and allows to document core microscopy concepts such as `<Light Source>`, `<Objective>, <Filter>, <Detector>`, etc.,), the OME specifications have not kept pace with the wide range of technologies that are now routinely used in the life and biomedical sciences.

#### 3.2.2 The 4DN-BINA proposal: a suite of three extensions of the core OME ontology

Due to its status as the *de facto* exchange specifications for imaging experiments, the robustness of its design, and the solid path forward toward modernization (Figure 4B and Text Box 1), the OME Data Model (i.e., OME core ontology) represents the ideal starting point for the suite of 4DN-BINA extensions presented here (Figure 5). As such, the 4DN-BINA-OME specifications proposal consists of three extensions of the OME Core (*4, 5*) that incorporate the concept of graded documentation requirements based on a tiered-system of guidelines (Figure 3; Table II). To achieve this goal, the 4DN-BINA-OME Microscopy Metadata Specifications (*7, 26, 73*) extend the core OME elements `<Instrument>` (Figure 5A) and `<Image>` (Figure 5B) to reflect the technological advances and the Quality Control requirements associated with state-of-the-art transmitted light, widefield- and confocal-fluorescence microscopy. Specifically:

1. The **<u>Basic extension</u>** is designed to better capture the technical complexity of transmitted light microscopy and wide-field fluorescence, including sub-pixel single-molecule localization and single-particle tracking experiments (Figure 5, blue and grey elements).
2. The **<u>Advanced and Confocal extension</u>** is designed to better capture experiments requiring tunable optics and confocal microscopy (Figure 5, green elements).
3. The **<u>Calibration and Performance extension</u>** introduces specifications for the capture of metrics required for microscope calibration and quantitative instrument performance assessment (Figure 5, maroon elements).

In order to facilitate understanding of the 4DN-BINA-OME by all relevant members of the community regardless of their information science expertise, while at the same time ensuring machine readability, formal representations of the 4DN-BINA-OME extensions are maintained on GitHub (*7*) in three formats: (1) a set of graphical ER schemas is used for facilitating an overall understanding of the model structure (*73*); (2) an excel spreadsheet expresses the details of the model in a human-readable form (*73*); finally (3) XML Schema Definition (XSD) is used to represent the model schema in a machine-readable manner (*26*).

##### 3.2.2.1 Basic 4DN-BINA-OME extension

The Basic 4DN-BINA extension of the OME Data Model was designed to better capture the technical complexity of transmitted light and wide-field fluorescence microscopy and is graphically presented in Figure 5, and Supplemental Figures 1A and 2 (*7, 26, 73*). This extension puts forth several types of modifications:

1. Extension of already existing elements such as `<Microscope>`, `<Laser>` `,<Objective>`, and `<Filter>` by the introduction of additional attributes (Figures 5, 6 and Supplemental Figure 1, blue and blue/red elements).
2. Introduction of novel elements such as `<StageInsert>, <SamplePositioning>, <FocusStabilizationDevice>` to capture the complexity of microscope hardware components commonly encountered in the field (Figures 5, 6 and Supplemental Figure 1, grey elements), and which combine with the new `<PlaneTransformMatrix>` affine transform element to encode locations in real-world coordinates
3. Mimicking the hierarchical structure of `<LightSource>`, introduction of several additional Abstract Parent Elements (APE) to describe hardware components, such as `<LightSourceCoupling>`, `<Filter>`, `<MirroringDevice>`, and `<Detector>`, that can be subdivided in specialized categories to streamline the structure of the model and avoid data duplication (i.e., the `<Detector>` category can be subdivided in `<Camera>` and `<PointDetector>)`.
4. Establishment of the concept of individual `<WavelengthRange>` to facilitate the description of multi-pass excitation sources, filters, dichroic-mirrors and detectors.
5. Introduction of additional concepts to better capture the settings of individual hardware components that are employed during the acquisition of a specific image and are stored in the core OME `<Image>` element (i.e.,`<MicroscopeSettings>, <CameraSettings> and <PMTSettings>`).
6. Expansion of the concept of `<LightPath>` to better capture the complexities of the light path that are typically encountered in different modern light microscopy modalities and uses a new `<LightPathMap>` element to describe the order of optical components that might be placed between the excitation source and the detector other than the filter and dichroic such as for example, `<LightSourceCoupling>, <Prism>, <PolarizationOptics>,<Lens>, and <OpticalAperture>`.
7. Introduction of the `<AdditionalDimensionMap>` and associated elements to flexibly handle dimensions such as fluorescence lifetime, polarization angle, and lambda beyond the five canonical X, Y, Z, T (time), and C (color) dimensions.

##### 3.2.2.2 Advanced and Confocal 4DN-BINA-OME Extension

This extension is designed to better capture experiments requiring the use of tunable optics and confocal microscopy (Figure 5, green elements). As depicted graphically in Figure 5, and Supplemental Figures 1B and 3 (*7, 26, 73*), this extension consists primarily in the introduction of novel concepts required to capture hardware components and image acquisition settings that are needed for confocal microscopy and other advanced acquisition modalities that require tunable excitation and emission light selection such as for example, `<ConfocalScanner>, <AcoustoOpticalBeamSplitter>,<LiquidCrystalTunableFilter>, and <PinHole>`.

##### 3.2.2.3 Calibration and Performance 4DN-BINA-OME Extension

The specifications for the capture of metrics required for light microscope calibration and quality control captured in this extension were developed in collaboration with QUAREP-LiMi (quarep.org; Figure 5, and Supplemental Figures 1C and 4) (*8, 9, 26, 37, 73*) and are described in detail in an accompanying manuscript (*21*). A diverse set of metrics (*41*–*51, 74*– *86*) can be used to measure microscope performance and control image quality depending on the type of experiment being performed and the questions being asked. Together, these measurements increase the depth and reliability of a variety of assessments, analyses, and comparisons performed on light microscopy images. Such calibration metrics can be subdivided into four categories: 1) optical; 2) intensity/excitation; 3) intensity/detector, and 4) mechanical (see also Table III in: Huisman et al., 2021) (*21*). Metrics in the first three categories evaluate the great majority of image data. Mechanical calibration metrics become most useful in experiments that involve time-lapse imaging or the tiling of multiple Fields Of View (FOV). In order to capture these metrics categories, the Calibration and Performance extension introduces the following new elements:

1. `<IntensityCalibrationTool>` and `<LightSensor>` represent hardware tools (e.g., power meter) used for performing specific intensity calibration procedures and represented as sub-elements of `<Instrument>`.
2. `<OpticalCalibration>`, `<ExcitationCalibration>, <DetectorCalibration>` and `<MechanicalCalibration>` store information that describes each respective calibration procedure, which might be performed to document individual image datasets and the resulting metrics. `<OpticalCalibration>`, and `<DetectorCalibration>` are connected with the `<Image>` element. In turn, `<ExcitationCalibration>` and `<MechanicalCalibration>` are associated with the `<Channel>` and the `<Plane>` elements respectively.
3. `<CalibrationStandardSlide>`, `<ColoredBeadsSlide>`, `<DNAOrigami>`, and`<FluorescenceReferenceSlide>,` which belong to the `<OpticalStandard>` group and store information that describe reference standards used for `<OpticalCalibration>` and other procedures including `<ChromaticRegistrationEvaluation>` and `<FieldUniformityEvaluation>.`

### 3.3 The third axis: model elements and attributes requirement levels

Along the third axis (Figure 6), individual metadata fields in these specifications are classified based on requirement level as described by the Request for Comment (RFC) document 2119 (*87*). The keyword MUST, or the terms “REQUIRED” or “SHALL,” mean that the definition is an absolute requirement to <u>validate experimental claims and ensure reproducibility</u>. The keyword SHOULD, or the adjective “RECOMMENDED,” mean that while there may exist valid reasons in particular circumstances to ignore a particular field, they are highly recommended to <u>maximize *Image Quality, scientific value and* FAIRness</u> (*22*). Two examples of the use of the third dimension to add flexibility to the proposed 4DN-BINA-OME Microscopy Metadata specifications are presented below:

<u>Example 1)</u> OME Core and 4DN-BINA Basic extension element `<Objective>` (Figure 6).

While the Manufacturer, Model, Magnification and Numerical Aperture (*LensNA*) of an objective are required to be able to interpret microscopy results and for reproducibility, other attributes such as a hardware component’s Lot Number, a Lens’s Back Focal Length and the Calibrated Magnification of an Objective are recommended to maximize Image Quality and scientific value but they are not required because they are not essential for reproducing the experiment.

<u>Example 2)</u> 4DN-BINA Calibration and Performance extension element `<ColorBeads>`

When using multicolored beads to prepare a colored beads slide to use for the optical calibration of a microscope, the Manufacturer, Catalog Number, and Concentration of the beads preparation alongside the Diameter of the beads are essential for the interpretation of the calibration results and for reproducibility. However, the bead’s Type and Material may be omitted because it can be argued that while that information improves the completeness of the data, they are not absolutely required for the correct interpretation of the results of the Optical calibration procedure in which the beads are utilized.

### 3.4 Model implementation: Material and Methods recommendations

A recent exploration about the quality of published Method sections in scientific articles containing images obtained with advanced microscopes, found that the quality of reporting was poor, with some articles containing no information about how images were obtained, and many articles lacking important basic details (*13*). Nonetheless, there is ample evidence that the publication of full details about how each image was obtained is vital for rigor, reproducibility and maximal scientific and sharing value (*14*–*16, 88*–*90*). In this context, the 4DN-BINA-OME Microscopy Metadata specifications presented are intended to provide a major contribution towards the development of community-driven criteria for which information should be included in the Methods sections of scientific publications.

As a first step, in close agreement with the proposal presented in parallel efforts (*89*)^23^, we propose that Microscopy Metadata appropriate for Tier 1 should always be included in the Material and Methods section of any journal publication to meet minimal rigor and reproducibility criteria (*13*). As such, the generalized and automated availability of Tier 1 metadata could save considerable effort both for authors, who would not need to search for information scattered across different data-files, hardware setups and lab notebooks in preparation for publication, and for readers, who would not need to search the various sections of publications for information that may or may not have been included.

### 3.5 Model implementation: Facilitated metadata collection

The importance of rich metadata to ensure the quality, reproducibility, as well as scientific and sharing value of image data cannot be overstated. However, the collection of rich sets of microscopy metadata is time-consuming and, in the absence of active participation from hardware manufacturers, imposes an unfair burden on experimental scientists and is therefore difficult to enforce. Appropriate community-validated software tools and data management practices are essential to streamline and automate the documentation of microscopy experiments. In this context, in parallel with this proposal for Microscopy Metadata guidelines a suite of three complementary and interoperable software tools are being developed and are presented in related manuscripts. 1) OMERO.mde (*28*) focuses on facilitating the consistent handling of image metadata ahead of data publication and deposition based on shared community Microscopy Metadata specifications and according to the FAIR principles. In addition, OMERO.mde promotes the early development of Image Metadata extension specifications to allow testing and validation before incorporation in community-accepted standards. 2) Micro-Meta App (*30, 91*) focuses on an easy-to-use, Graphical User Interface (GUI)-based platform that interactively guides users through the process of building tier-based records of microscope hardware, accessories and image acquisition settings containing all relevant Microscopy Metadata as sanctioned by the community specifications such as the ones described here. Because of its graphical nature, Micro-Meta App is particularly suited for imaging scientists to enter all microscope metadata and use the tool for teaching trainees about Microscopy, and training microscope-users in imaging facilities. 3) Finally, MethodsJ2 ^(*29*)^ focuses on automating the process of writing Microscopy Metadata guidelines-compliant Methods and Acknowledgment sections for scientific publications utilizing microscopy experiments. MethodsJ2, by design, operates in concert to automatically import Microscopy Metadata from the Micro-Meta App (*30, 91*).

### 3.6 Model implementation: Information required for basic image interpretability

To ensure the basic interpretability of image data acquired before the adoption of community-sanctioned guidelines, any data that might be shared or published should, at the very least, contain the required metadata fields stipulated by the intersection between Tier 1 and the Core of the OME Data Model. Thus, Tier 1/Core sanctions the baseline metadata requirements for any light microscopy experiment to be interpretable, utilized and shared for scientific purposes. Specifically, this includes minimal microscope Hardware Specifications (i.e., microscope, light source and objective manufacturer information and essential description), and essential information about the Image structure (i.e., number of planes, channels and time-points, pixel size, fluorophore name, emission and excitation wavelength, etc.).

## 4 - CONCLUSION

Light Microscopy images need to be accompanied by thorough documentation of the microscope hardware and imaging settings to ensure a correct interpretation of the results. A significant challenge with the reproducibility of microscopy results and their integration with other data types, such as chromatin folding maps generated by the 4DN consortium (*1, 2*), lies in the lack of standardized reporting guidelines for microscopy experiments as well as instrument performance and calibration standards (*13*–*15, 88*). Despite a growing consensus that such standards for light microscopy are desirable, previous efforts to develop shared microscopy data models and application programming interfaces (*4, 5, 54*) have not yet succeeded in the establishment of a universal set of norms. In this manuscript, a framework to extend the OME Data Model is put forth to help address this challenge. In addition to aligning the OME Data Model to current technological developments, the specifications advanced here focus on the maximization of usability via the introduction of a tiered system of documentation requirements, on an expandable suite of model extensions, including the first available data model for quality-control metadata for light microscopy imaging and flexible use of required, and recommended, fields.

Microscopy is not the only field in which recent technological advances have resulted in increasingly rich datasets. Recent examples are genomic DNA and transcriptomics RNA sequencing, which are, in fact, much younger fields than microscopy. While protocols varied substantially in their early days (the original images from the sequencer were kept with the determined sequence), it took only about a decade to establish metadata requirements. One factor that helped establish such metadata criteria was the NIH Encyclopedia of DNA Elements (ENCODE) consortium (*10, 11*). The development of Standard Operating Procedures (SOPs) and shared benchmarks (i.e., gold-standards) within this group was pivotal for the establishment of agreeable standards for practical day-to-day use. In the interest of scientific progress and making data FAIR, data and metadata standards should not be dictated by individual laboratories or microscope manufacturers. Instead, they should emerge organically from discussions involving all members of the community who can benefit from standardization and be subjected to evaluation before adoption.

In this spirit, the initial draft Microscopy Metadata specifications put forth by the 4DN (*1, 2*) IWG were evaluated and revised by the BINA QC-DM-WG (*3*), resulting in the current proposal. Furthermore, this process is being carried out in alliance with the QUAREP-LiMi initiative ((*9, 37*) to ensure that all participating imaging community stakeholders (importantly including microscope and software tool manufacturers, who are ultimately responsible for providing the information to be recorded in microscopy metadata) are involved from the ground up and provide timely feedback. Because it is inherently impossible to predict all future changes the light microscopy field might undergo and in order to ensure rigor and reproducibility for image data now and in the future, it is essential that the 4DN-BINA (as well as future) extensions of the OME Data Model for bioimaging metadata proposed here are capable of gradually evolving to capture any future technical development while supporting FAIR data principles(*22, 28*). This is particularly important in the face of the establishment of a growing number of public image data resources (*20*) such as the IDR (*55*), EMPIAR (*59*), and Bioimage Archive (*60*) hosted at the EMBL - EBI; the Japanese SSBD hosted by RIKEN (*64*); and, in the USA, the NIH CELL Image Library (*65*) the BRAIN initiative’s brain tissue imaging resources^17^. These resources offer the opportunity to emulate for light microscopy the successful path that has led to community standards in the field of genomics (*92*–*96*). To this aim, in addition to our work in the context of QUAREP-LiMi, the further development of the Microscopy Metadata Specifications is being coordinated with other parallel initiatives including: 1) the OME community development of general criteria and procedures to capture and store metadata in OME-NGFF (Text Box 1). The OME NGFF effort (*35, 36*) is implementing storage approaches to hold the binary pixel data and the metadata described herein in standardized, shareable, long-lived, efficient, and performant containers (e.g. files). 2) The EMBL-EBI development of the REMBI recommendations for metadata to be included with imaging datasets deposited to BioImage Archive (*60, 66, 97*). 3) The development of the International Standards Organization (ISO) 23494-1 standard that will include the 4D**N**-**B**INA-**O**ME (NBO) Microscopy Metadata specifications as part of a Provenance information model for biological material and data (*34, 98*). 4) The development of educational material in collaboration with Global Bioimaging to increase awareness about the importance of metadata standards to ensure image data quality, reproducibility and re-use value.

In conclusion, we are convinced that because of its strong roots in the community, and because it is closely linked with the parallel development of easy-to-use interactive tools to facilitate metadata collection (*28*–*30*), the flexible model framework presented here will provide a significant step forward towards the establishment of robust and future-proof light microscopy metadata standards. In turn, this will help increase rigor and reproducibility in imaging data, rewarding everyone involved with improved trust in published results.

## Abbreviation list

OME: Open Microscopy Environment

## 5 – AUTHORS CONTRIBUTIONS

Author contributions categories utilized here were devised by the CRediT initiative (*99, 100*).

**Mathias Hammer**: Conceptualization, Methodology, Investigation, Data Curation, Writing - Review & Editing; **Maximiliaan Huisman**: Conceptualization, Methodology, Investigation, Writing - Original Draft, Writing - Review & Editing, Visualization; **Alessandro Rigano:** Conceptualization, Methodology, Software; **Ulrike Boehm, James J. Chambers, Nathalie Gaudreault, Alison J. North, Jaime A. Pimentel**, and **Damir Sudar**: Validation, Investigation, Data Curation, Writing - Review & Editing; **Peter Bajcsy, Claire M. Brown, Alexander D. Corbett, Orestis Faklaris, Judith Lacoste, Alex Laude, Glyn Nelson**, and **Roland Nitschke**: Validation, Writing - Review & Editing; **Farzin Farzam**: Investigation; **Carlas Smith**: Conceptualization, Methodology; **David Grunwald**: Conceptualization, Methodology, Investigation, Resources, Writing - Original Draft, Supervision, Project administration, Funding acquisition; **Caterina Strambio-De-Castillia**: Conceptualization, Methodology, Software, Validation, Investigation, Resources, Data Curation, Writing - Original Draft, Writing - Review & Editing, Visualization, Supervision, Project administration, Funding acquisition.

## 6 - ACKNOWLEDGEMENTS

We would like to thank Kevin Fogarty, Lawrence Lifshitz and Karl Bellve at the Biomedical Imaging Group of the Program in Molecular Medicine at the University of Massachusetts Medical School for invaluable intellectual input and countless fruitful discussions and for their friendship, advice, and steadfast support throughout the development of this project.

This project could never have been carried out without the leadership, insightful discussions, support and friendship of all OME consortium members with particular reference to Jason Swedlow, Josh Moore, Chris Allan, Jean Marie Burel, and Will Moore. We are massively indebted to the RIKEN community for their fantastic work to bring open science into biology. We would like to particularly acknowledge Norio Kobayashi and Shuichi Onami for their friendship and support.

We thank all members of BioImaging North America (in particular Lisa Cameron, Michelle Itano, and Paula Montero-Llopis), German Bioimaging (in particular Susanne Kunis and Stephanie Wiedkamp Peters), Euro-Bioimaging (in particular Antje Keppler and Federica Paina) and QUAREP-LiMi (in particular all members of the Working Group 7 - Metadata; quarep.org) for invaluable intellectual input, fruitful discussions and advice. We are also indebted to the following individuals for their continued and steadfast support: Jeremy Luban, Roger Davis, and Thoru Pederson at the University of Massachusetts Medical School; Burak Alver, Joan Ritland, Rob Singer, and Warren Zipfel at the 4D Nucleome Project; Ian Fingerman, John Satterlee, Judy Mietz, Richard Conroy, and Olivier Blondel at the NIH. We would like to thank Dr. Darryl Conte for the critical reading of the manuscript. We are deeply indebted to Thao P. Do, Cell visualization and web specialist at the Allen Institute for Cell Science, for her expert advice and skilled work, which was invaluable for the production of Figures 1, 2, 3 and 4.

This work was supported by NIH grant #1U01EB021238 and NSF grant #1917206 to D.G., NIH grant # 2U01CA200059-06 to C.S.D.C and D.G., and by grant #2019-198155 (5022) awarded to C.S.D.C. by the Chan Zuckerberg Initiative DAF, an advised fund of Silicon Valley Community Foundation, as part of their Imaging Scientist Program. D.S. was funded in part by NIH/NCI grants U54CA209988 and U2CCA23380. C.M.B. was funded in part by grant #2020-225398 from the Chan Zuckerberg Initiative DAF, an advised fund of Silicon Valley Community Foundation. R.N. was funded by the Deutsche Forschungsgemeinschaft (DFG, German Research Foundation) grant number Ni 451/9-1 MIAP-Freiburg. C.S.S. was supported by the Netherlands Organisation for Scientific Research (NWO), under NWO START-UP project no. 740.018.015 and NWO Veni project no. 16761.

## TEXT BOXES

### Text Box I

#### Charting a solid path towards next-generation storage mechanisms for community-driven, OME-based Microscopy Image Data Standards

Microscopy Metadata, stored (Figure 1, ***HOW*** *yellow* and ***WHERE*** *blue bubbles*) following the Open Microscopy Environment (OME) Data Model(*4, 5*) is represented in the form of OME-Extensible Markup Language (OME-XML), which is typically stored in the header of OME-TIFF files. Consequently, the XML Schema Definition (XSD) formalism is used to represent the model schema in a machine-readable manner. However, despite its advantages, XSD is not ideally suited to allow the OME Data Model to serve as the foundation for the community-development and maintenance of globally accepted light microscopy standards (Figure 1). Because XSD does not support the storage of novel types of information within the core of the model, the capture of ever-evolving microscopy technologies and modalities requires the periodical release of new versions of the OME XSD schema (*6*) accompanied by XML Stylesheet Language (XSL) based templates for making sure legacy documents could be kept up to date. This burdensome process is ultimately unsustainable. Consequently, it is necessary to develop new strategies with a more open paradigm.

Under this new paradigm, one would assume that no single authority exists to decide which information must be recorded in metadata models making it necessary for commonly used concepts to be incorporated over time into community-driven standards. In this context, agreement has to be reached not much on ***WHAT*** concepts have to be recorded for the documentation of imaging experiments (Figure 1, *magenta bubble*), rather on the development of shared mechanisms defining ***HOW*** new types of (meta)data have to be recorded (Figure 1, *yellow bubble*) and associated with the Image data file format (Figure 1, ***WHERE*** *blue bubble*) (*35*).

In this context, the OME consortium, in collaboration with RIKEN, has started experimenting with the idea of utilizing Resource Description Framework (RDF) (*101, 102*) triples conforming to the Web Ontology Language (OWL) (*103*) to describe OME-compatible image metadata (*32, 33, 104, 105*) and be incorporated in the Next-Generation File Formats (NGFF) currently being developed by the OME consortium (*35*). By employing this method, it would be possible for users to produce, find and access quality-controlled image data for re-analysis and integration. Specifically, the depicted method will provide two major advantages:

1. Individual groups specializing in different aspects of the imaging world will have equal status and a shared path to develop new areas of the model (*28*). In turn, this will provide a method for different communities to collectively develop a complete picture (Figure 2) of all the information required to ensure rigor and reproducibility for modern imaging experiments.
2. At the same time, community-driven standards could evolve gradually over time by incorporating novel concepts into the core as they are developed peripherally from the core, vetted by the community, and commonly adopted.

As a *proof-of-concep*t, an implementation of the OME Data Model was built in RDF/OWL(*106*), and applied to the modeling of specifications defined by the 4D Nucleome Imaging Standard Working Group(*1, 2*) for the exchange of image data and integration with genomics datasets (*31, 107*). This demonstrated the potential utility of this approach, laying the foundation for ongoing community discussions to identify the path of choice for modern Light Microscopy Image Data Standards (Figure 1).

## Footnotes

17 https://doryworkspace.org/

## Notes

https://zenodo.org/record/4710731

https://zenodo.org/record/4711229

https://zenodo.org/record/4711426

### Competing Interest Statement

The authors have declared no competing interest.

### Summary of Updates

A few small revisions: -Reference update -Minor revision in Table I

## REFERENCES

1. 4D Nucleome Consortium, The 4D Nucleome Web Portal. 4dnucleome.org (2017), (available at https://www.4dnucleome.org/).

2. J. Dekker, A. S. Belmont, M. Guttman, V. O. Leshyk, J. T. Lis, S. Lomvardas, L. A. Mirny, C. C. O’Shea, P. J. Park, B. Ren, J. C. R. Politz, J. Shendure, S. Zhong, 4D Nucleome Network, The 4D nucleome project. Nature. 549, 219–226 (2017).

3. C. Strambio-De-Castillia, P. Bajcsy, U. Boehm, J. Chambers, A. D. Corbett, O. Faklaris, N. Gaudreault, J. Lacoste, A. Laude, G. Nelson, R. Nitschke, J. A. Pimentel, D. Sudar, C. M. Brown, A. J. North, Quality Control and Data Management | Bioimaging North America (BINA). Bioimaging North America (2019), (available at https://www.bioimagingna.org/qc-dm-wg).

4. M. Linkert, C. T. Rueden, C. Allan, J.-M. Burel, W. Moore, A. Patterson, B. Loranger, J. Moore, C. Neves, D. MacDonald, A. Tarkowska, C. Sticco, E. Hill, M. Rossner, K.W. Eliceiri, J.R. Swedlow, Metadata matters: access to image data in the real world. J. Cell Biol. 189, 777–782 (2010).

5. I. G. Goldberg, C. Allan, J.-M. Burel, D. Creager, A. Falconi, H. Hochheiser, J. Johnston, J. Mellen, P. K. Sorger, J. R. Swedlow, The Open Microscopy Environment (OME) Data Model and XML file: open tools for informatics and quantitative analysis in biological imaging. Genome Biol. 6, R47 (2005).

6. OME Consortium, “OME Data Model and File Formats 6.2.2 Documentation — OME Data Model and File Formats 6.2.2 documentation” (v6.2.2, openmicroscopy.org, 2016), (available at https://docs.openmicroscopy.org/ome-model/6.2.2/).

7. A. Rigano, U. Boehm, J. J. Chambers, N. Gaudreault, A. J. North, J. A. Pimentel, D. Sudar, P. Bajcsy, C. M. Brown, A. D. Corbett, O. Faklaris, J. Lacoste, A. Laude, G. Nelson, R. Nitschke, D. Grunwald, C. Strambio-De-Castillia, 4DN-BINA-OME (NBO) Tiered Microscopy Metadata Specifications - v2.01 (https://github.com/WU-BIMAC, 2021; https://zenodo.org/record/4710731).

8. M. Hammer, M. Huisman, A. Rigano, U. Boehm, J. J. Chambers, N. Gaudreault, J. A. Pimentel, D. Sudar, P. Bajcsy, C. M. Brown, A. D. Corbett, O. Faklaris, J. Lacoste, A. Laude, G. Nelson, R. Nitschke, A. J. North, R. Gopinathan, F. Farzam, C. Smith, W. Gipfel, J. Ritland, D. Grunwald, C. Strambio-De-Castillia, 4DN-BINA-OME (NBO)-Microscopy Metadata Specifications - Tiers System_v2.01 (GitHub - https://github.com/WU-BIMAC/NBOMicroscopyMetadataSpecs, 2021; https://github.com/WU-BIMAC/NBOMicroscopyMetadataSpecs/tree/master/Tier%20System/stable%20version/v02-01).

9. G. Nelson, U. Boehm, S. Bagley, P. Bajcsy, J. Bischof, C. M. Brown, A. Dauphin, I. M. Dobbie, J. E. Eriksson, O. Faklaris, J. Fernandez-Rodriguez, A. Ferrand, L. Gelman, A. Gheisari, H. Hartmann, C. Kukat, A. Laude, M. Mitkovski, S. Munck, A. J. North, T. M. Rasse, U. Resch-Genger, L. C. Schuetz, A. Seitz, C. Strambio-De-Castillia, J. R. Swedlow, I. Alexopoulos, K. Aumayr, S. Avilov, G.-J. Bakker, R. R. Bammann, A. Bassi, H. Beckert, S. Beer, Y. Belyaev, J. Bierwagen, K. A. Birngruber, M. Bosch, J. Breitlow, L. A. Cameron, J. Chalfoun, J. J. Chambers, C.-L. Chen, E. Conde-Sousa, A. D. Corbett, F. P. Cordelieres, E. Del Nery, R. Dietzel, F. Eismann, E. Fazeli, A. Felscher, H. Fried, N. Gaudreault, W. I. Goh, T. Guilbert, R. Hadleigh, P. Hemmerich, G. A. Holst, M. S. Itano, C. B. Jaffe, H. K. Jambor, S. C. Jarvis, A. Keppler, D. Kirchenbuechler, M. Kirchner, N. Kobayashi, G. Krens, S. Kunis, J. Lacoste, M. Marcello, G. G. Martins, D. J. Metcalf, C. A. Mitchell, J. Moore, T. Mueller, M. S. Nelson, S. Ogg, S. Onami, A. L. Palmer, P. Paul-Gilloteaux, J. A. Pimentel, L. Plantard, S. Podder, E. Rexhepaj, M. Royeck, A. Royon, M. A. Saari, D. Schapman, V. Schoonderwoert, B. Schroth-Diez, S. Schwartz, M. Shaw, M. Spitaler, M. T. Stoeckl, D. Sudar, J. Teillon, S. Terjung, R. Thuenauer, C. D. Wilms, G. D. Wright, R. Nitschke, QUAREP-LiMi: A community-driven initiative to establish guidelines for quality assessment and reproducibility for instruments and images in light microscopy. arXiv [q-bio.OT] (2021), (available at http://arxiv.org/abs/2101.09153).

10. ENCODE Project Consortium, An integrated encyclopedia of DNA elements in the human genome. Nature. 489, 57–74 (2012).

11. N. de Souza, The ENCODE project. Nat. Methods. 9 (2012), p. 1046.

12. M. Baker, D. Penny, 1,500 scientists lift the lid on reproducibility. Nature. 533 (2016), pp. 452–454.

13. G. Marqués, T. Pengo, M. A. Sanders, Imaging methods are vastly underreported in biomedical research. Elife. 9 (2020), doi:10.7554/eLife.55133.

14. M. R. Sheen, J. L. Fields, B. Northan, J. Lacoste, L.-H. Ang, S. Fiering, Reproducibility Project: Cancer Biology, Replication Study: Biomechanical remodeling of the microenvironment by stromal caveolin-1 favors tumor invasion and metastasis. Elife. 8 (2019),, doi:10.7554/eLife.45120.

15. M. P. Viana, J. Chen, T. A. Knijnenburg, R. Vasan, C. Yan, J. E. Arakaki, M. Bailey, B. Berry, A. Borensztejn, J. M. Brown, S. Carlson, J. A. Cass, B. Chaudhuri, K. R. Cordes Metzler, M. E. Coston, Z. J. Crabtree, S. Davidson, C. M. DeLizo, S. Dhaka, S. Q. Dinh, T. P. Do, J. Domingus, R. M. Donovan-Maiye, T. J. Foster, C. L. Frick, G. Fujioka, M. A. Fuqua, J. L. Gehring, K. A. Gerbin, T. Grancharova, B. W. Gregor, L. J. Harrylock, A. Haupt, M. C. Hendershott, C. Hookway, A. R. Horwitz, C. Hughes, E. J. Isaac, G. R. Johnson, B. Kim, A. N. Leonard, W. W. Leung, J. J. Lucas, S. A. Ludmann, B. M. Lyons, H. Malik, R. McGregor, G. E. Medrash, S. L. Meharry, K. Mitcham, I. A. Mueller, T. L. Murphy-Stevens, A. Nath, A. M. Nelson, L. Paleologu, T. Alexander Popiel, M. M. Riel-Mehan, B. Roberts, L. M. Schaefbauer, M. Schwarzl, J. Sherman, S. Slaton, M. Filip Sluzewski, J. E. Smith, Y. Sul, M. J. Swain-Bowden, W. Joyce Tang, D. J. Thirstrup, D. M. Toloudis, A. P. Tucker, V. Valencia, W. Wiegraebe, T. Wijeratna, R. Yang, R. J. Zaunbrecher, Allen Institute for Cell Science, G. T. Johnson, R. N. Gunawardane, N. Gaudreault, J. A. Theriot, S. M. Rafelski, Robust integrated intracellular organization of the human iPS cell: where, how much, and how variable. BioRxiv.org (2021), p. 2020.12.08.415562.

16. R. Botvinik-Nezer, F. Holzmeister, C. F. Camerer, A. Dreber, J. Huber, M. Johannesson, M. Kirchler, R. Iwanir, J. A. Mumford, R. A. Adcock, P. Avesani, B. M. Baczkowski, A. Bajracharya, L. Bakst, S. Ball, M. Barilari, N. Bault, D. Beaton, J. Beitner, R. G. Benoit, R. M. W. J. Berkers, J. P. Bhanji, B. B. Biswal, S. Bobadilla-Suarez, T. Bortolini, K. L. Bottenhorn, A. Bowring, S. Braem, H. R. Brooks, E. G. Brudner, C. B. Calderon, J. A. Camilleri, J. J. Castrellon, L. Cecchetti, E. C. Cieslik, Z. J. Cole, O. Collignon, R. W. Cox, W. A. Cunningham, S. Czoschke, K. Dadi, C. P. Davis, A. D. Luca, M. R. Delgado, L. Demetriou, J. B. Dennison, X. Di, E. W. Dickie, E. Dobryakova, C. L. Donnat, J. Dukart, N. W. Duncan, J. Durnez, A. Eed, S. B. Eickhoff, A. Erhart, L. Fontanesi, G. M. Fricke, S. Fu, A. Galván, R. Gau, S. Genon, T. Glatard, E. Glerean, J. J. Goeman, S. A. E. Golowin, C. González-García, K. J. Gorgolewski, C. L. Grady, M. A. Green, J. F. Guassi Moreira, O. Guest, S. Hakimi, J. P. Hamilton, R. Hancock, G. Handjaras, B. B. Harry, C. Hawco, P. Herholz, G. Herman, S. Heunis, F. Hoffstaedter, J. Hogeveen, S. Holmes, C.P. Hu, S. A. Huettel, M. E. Hughes, V. Iacovella, A. D. Iordan, P. M. Isager, A. I. Isik, A. Jahn, M. R. Johnson, T. Johnstone, M. J. E. Joseph, A. C. Juliano, J. W. Kable, M. Kassinopoulos, C. Koba, X.-Z. Kong, T. R. Koscik, N. E. Kucukboyaci, B. A. Kuhl, S. Kupek, A. R. Laird, C. Lamm, R. Langner, N. Lauharatanahirun, H. Lee, S. Lee, A. Leemans, A. Leo, E. Lesage, F. Li, M. Y. C. Li, P. C. Lim, E. N. Lintz, S. W. Liphardt, A. B. Losecaat Vermeer, B. C. Love, M. L. Mack, N. Malpica, T. Marins, C. Maumet, K. McDonald, J. T. McGuire, H. Melero, A.S. Méndez Leal, B. Meyer, K. N. Meyer, G. Mihai, G. D. Mitsis, J. Moll, D. M. Nielson, G. Nilsonne, M. P. Notter, E. Olivetti, A. I. Onicas, P. Papale, K. R. Patil, J. E. Peelle, A. Pérez, D. Pischedda, J.B. Poline, Y. Prystauka, S. Ray, P. A. Reuter-Lorenz, R. C. Reynolds, E. Ricciardi, J. R. Rieck, A. M. Rodriguez-Thompson, A. Romyn, T. Salo, G. R. Samanez-Larkin, E. Sanz-Morales, M. L. Schlichting, D. H. Schultz, Q. Shen, M. A. Sheridan, J. A. Silvers, K. Skagerlund, A. Smith, D. V. Smith, P. Sokol-Hessner, S. R. Steinkamp, S. M. Tashjian, B. Thirion, J. N. Thorp, G. Tinghög, L. Tisdall, S. H. Tompson, C. Toro-Serey, J. J. Torre Tresols, L. Tozzi, V. Truong, L. Turella, A. E. van ‘t Veer, T. Verguts, J. M. Vettel, S. Vijayarajah, K. Vo, M. B. Wall, W. D. Weeda, S. Weis, D. J. White, D. Wisniewski, A. Xifra-Porxas, E. Yearling, S. Yoon, R. Yuan, K. S. L. Yuen, L. Zhang, X. Zhang, J. E. Zosky, T. E. Nichols, R. A. Poldrack, T. Schonberg, Variability in the analysis of a single neuroimaging dataset by many teams. Nature. 582, 84–88 (2020).

17. Nature Editorial Staff, Better research through metrology. Nat. Methods. 15, 395 (2018).

18. J. Pines, Image integrity and standards. Open Biol. 10 (2020), p. 200165.

19. J. Eriksson, I. Pukonen, “D2.3 Common international recommendation for quality assurance and management in open access imaging infrastructures” (Global BioImaging Project, 2018), (available at https://www.globalbioimaging.org/user/pages/05.documents/D2.3_Publication%20of%20common%20recommendation_quality%20assurance%20and%20management%20in%20open%20access%20imaging%20infrastructures.pdf).

20. J. R. Swedlow, P. Kankaanpää, U. Sarkans, W. Goscinski, G. Galloway, R. P. Sullivan, C. M. Brown, C. Wood, A. Keppler, B. Loos, S. Zullino, D. L. Longo, S. Aime, S. Onami, A Global View of Standards for Open Image Data Formats and Repositories. arXiv [q-bio.OT] (2020), (available at http://arxiv.org/abs/2010.10107).

21. M. Huisman, M. Hammer, A. Rigano, U. Boehm, J. J. Chambers, N. Gaudreault, J. A. Pimentel, D. Sudar, P. Bajcsy, C. M. Brown, A. D. Corbett, O. Faklaris, J. Lacoste, A. Laude, G. Nelson, R. Nitschke, A. J. North, D. Grunwald, C. Strambio-De-Castillia, A perspective on Microscopy Metadata: data provenance and quality control. arXiv [q-bio.QM] (2021), (available at https://arxiv.org/abs/1910.11370).

22. M. D. Wilkinson, M. Dumontier, I. J. J. Aalbersberg, G. Appleton, M. Axton, A. Baak, N. Blomberg, J.-W. Boiten, L. B. da Silva Santos, P. E. Bourne, J. Bouwman, A. J. Brookes, T. Clark, M. Crosas, I. Dillo, O. Dumon, S. Edmunds, C. T. Evelo, R. Finkers Gonzalez-Beltran, A. J. G. Gray, P. Groth, C. Goble, J. S. Grethe, J. Heringa, P. A. C. ‘t Hoen, R. Hooft, T. Kuhn, R. Kok, J. Kok, S. J. Lusher, M. E. Martone, A. Mons, A. L. Packer, B. Persson, P. Rocca-Serra, M. Roos, R. van Schaik, S.-A. Sansone, E. Schultes, T. Sengstag, T. Slater, G. Strawn, M. A. Swertz, M. Thompson, J. van der Lei, E. van Mulligen, J. Velterop, A. Waagmeester, P. Wittenburg, K. Wolstencroft, J. Zhao, B. Mons, The FAIR Guiding Principles for scientific data management and stewardship. Sci Data. 3, 160018 (2016).

23. S. Li, S. Besson, C. Blackburn, M. Carroll, R. K. Ferguson, H. Flynn, K. Gillen, R. Leigh, D. Lindner, M. Linkert, W. J. Moore, Ramalingam, E. Rozbicki, G. Rustici, A. Tarkowska, P. Walczysko, E. Williams, C. Allan, J.-M. Burel, J. Moore, J. R. Swedlow, Metadata management for high content screening in OMERO. Methods. 96, 27–32 (2016).

24. J.-M. Burel, S. Besson, C. Blackburn, M. Carroll, R. K. Ferguson, H. Flynn, K. Gillen, R. Leigh, S. Li, D. Lindner, M. Linkert, W. J. Moore, B. Ramalingam, E. Rozbicki, A. Tarkowska, P. Walczysko, C. Allan, J. Moore, J. R. Swedlow, Publishing and sharing multi-dimensional image data with OMERO. Mamm. Genome. 26, 441–447 (2015).

25. S. Besson, R. Leigh, M. Linkert, C. Allan, J.-M. Burel, M. Carroll, D. Gault, R. Gozim, S. Li, D. Lindner, J. Moore, W. Moore, P. Walczysko, F. Wong, J. R. Swedlow, Bringing Open Data to Whole Slide Imaging. Digit Pathol (2019). 2019, 3–10 (2019).

26. A. Rigano, C. Strambio-De-Castillia, 4DN-BINA-OME (NBO) Tiered Microscopy Metadata Specifications - v2.01 - XSD schema (https://github.com/WU-BIMAC/NBOMicroscopyMetadataSpecs, 2021; https://zenodo.org/record/4711426).

27. D. P. W. Russell, P. K. Sorger, Maintaining the provenance of microscopy metadata using OMERO.forms software. BiorXiv, 109199 (2017).

28. S. Kunis, S. Hänsch, C. Schmidt, F. Wong, S. Weidtkamp-Peters, OMERO.mde in a use case for microscopy metadata harmonization: Facilitating FAIR principles in practical application with metadata annotation tools. arXiv [q-bio.QM] (2021), (available at http://arxiv.org/abs/2103.02942).

29. J. Ryan, T. Pengo, A. Rigano, P. Montero Llopis, M. S. Itano, L. C. Cameron, G. Marqués, C. Strambio-De-Castillia, M. A. Sanders, C. M. Brown, MethodsJ2: A Tool to Help Improve Microscopy Methods Reporting. Nat. Methods (Manuscript under joint submission; personal communication).

30. A. Rigano, S. Ehmsen, S. U. Ozturk, J. Ryan, A. Balashov, M. Hammer, K. Kirli, U. Boehm, C. M. Brown, K. Belve’, J. Chambers, A. Cosolo, R. Coleman, O. Faklaris, K. Fogarty, T. Guilbert, A. B. Hamacher, M. S. Itano, D. P. Keeley, S. Kunis, J. Lacoste, A. Laude, W. Ma, M. Marcello, P. Montero-Llopis, G. Nelson, R. Nitschke, J. A. Pimentel, S. Weidtkamp-Peters, P. Park, B. Alver, D. Grunwald, C. Strambio-De-Castillia, Micro-Meta App: an interactive software tool to facilitate the collection of microscopy metadata based on community-driven specifications. BioRxiv (2021), doi:https://www.biorxiv.org/content/10.1101/2021.05.31.446382v1.

31. M. Hammer, A. Rigano, F. Farzam, M. Huisman, D. Grünwald, C. Strambio-De-Castillia, in Proceedings of the 12th Annual SWAT4(HC)LS (Semantic Web Applications and Tools for Healthcare and Life Sciences) Conference, A. Burger, R. Cornet, A. Waagmeester, Eds. (figshare, 2020; http://dx.doi.org/10.6084/M9.FIGSHARE.12758957), p. 32.

32. N. Kobayashi, J. Moore, S. Onami, J. R. Swedlow, in Proceedings of the 12th SWAT4(HC)LS (Semantic Web Applications and Tools for Healthcare and Life Sciences) Conference, A. Burger, R. Cornet, A. Waagmeester, Eds. (http://ceur-ws.org, 2019), p. 29.

33. J. Moore, N. Kobayashi, S. Kunis, S. Onami, J. R. Swedlow, the OME Consortium, in Proceedings of the 12th SWAT4(HC)LS (Semantic Web Applications and Tools for Healthcare and Life Sciences) Conference, A. Burger, R. Cornet, A. Waagmeester, Eds. (http://ceur-ws.org, 2019), p. 17.

34. R. Wittner, P. Holub, J. Geiger, H. Müller, C. Goble, S. Soiland-Reyes, E. Fairweather, L. Pirredu, F. Frexia, C. Mascia, G. Zanetti, H. Nakae, C. Strambio-De-Castillia, J. Moore, D. Grunwald, J. Swedlow, ISO 23494: Biotechnology - provenance information model for biological specimen and data (2020),, doi:10.5281/ZENODO.3901011.

35. J. Moore, S. Besson, OME Consortium, Next-generation file format (NGFF) specifications for storing bioimaging data in the cloud (2021),, doi:10.5281/zenodo.4282107.

36. J. Moore, C. Allan, S. Besson, J.-M. Burel, E. Diel, D. Gault, K. Kozlowski, D. Lindner, M. Linkert, T. Manz, W. Moore, C. Tischer, J. R. Swedlow, OME-NGFF: scalable format strategies for interoperable bioimaging data. bioRxiv (2021), p. 2021.03.31.437929.

37. U. Boehm, G. Nelson, C. M. Brown, S. Bagley, P. Bajcsy, J. Bischof, A. Dauphin, I. M. Dobbie, J. E. Eriksson, O. Faklaris, J. Fernandez-Rodriguez, A. Ferrand, L. Gelman, A. Gheisari, H. Hartmann, C. Kukat, A. Laude, M. Mitkovski, S. Munck, A. J. North, T. M. Rasse, U. Resch-Genger, L. C. Schuetz, A. Seitz, C. Strambio-De-Castillia, J. R. Swedlow, R. Nitschke, QUAREP-LiMi: A community endeavour to advance Quality Assessment and Reproducibility in Light Microscopy. Nat. Methods. ePub (2021),, doi:10.1038/s41592-021-01162-y.

38. K. Miura, S.F. Nørrelykke, Reproducible image handling and analysis. EMBO J., e105889 (2021).

39. A. Rigano, C. Strambio-De-Castillia, Proposal for minimum information guidelines to report and reproduce results of particle tracking and motion analysis. bioRxiv, 155036 (2017).

40. P. Masuzzo, L. Huyck, A. Simiczyjew, C. Ampe, L. Martens, M. Van Troys, An end-to-end software solution for the analysis of high-throughput single-cell migration data. Sci. Rep. 7, 42383 (2017).

41. A. J. North, Seeing is believing? A beginners’ guide to practical pitfalls in image acquisition. J. Cell Biol. 172, 9–18 (2006).

42. J. Demmerle, C. Innocent, A. J. North, G. Ball, M. Müller, E. Miron, A. Matsuda, I. M. Dobbie, Y. Markaki, L. Schermelleh, Strategic and practical guidelines for successful structured illumination microscopy. Nat. Protoc. 12, 988–1010 (2017).

43. J. M. Murray, P. L. Appleton, J. R. Swedlow, J. C. Waters, Evaluating performance in three-dimensional fluorescence microscopy. J. Microsc. 228, 390–405 (2007).

44. J. C. Waters, Accuracy and precision in quantitative fluorescence microscopy. J. Cell Biol. 185, 1135–1148 (2009).

45. T. J. Lambert, J. C. Waters, Assessing camera performance for quantitative microscopy. Methods Cell Biol. 123, 35–53 (2014).

46. L. J. Petrak, J. C. Waters, A practical guide to microscope care and maintenance. Methods Cell Biol. 123, 55–76 (2014).

47. A. P.-T. Jost, J. C. Waters, Designing a rigorous microscopy experiment: Validating methods and avoiding bias. J. Cell Biol. 218, 1452–1466 (2019).

48. R. C. Deagle, T.-L. E. Wee, C. M. Brown, Reproducibility in light microscopy: Maintenance, standards and SOPs. Int. J. Biochem. Cell Biol. 89, 120–124 (2017).

49. F. Mubaid, D. Kaufman, T.-L. Wee, D.-S. Nguyen-Huu, D. Young, M. Anghelopoulou, C. M. Brown, Fluorescence microscope light source stability. Histochem. Cell Biol. 151, 357–366 (2019).

50. P. Theer, C. Mongis, M. Knop, PSFj: know your fluorescence microscope. Nat. Methods. 11, 981–982 (2014).

51. T. A. Ferreira, A. V. Blackman, J. Oyrer, S. Jayabal, A. J. Chung, A. J. Watt, P.J. Sjöström, D. J. van Meyel, Neuronal morphometry directly from bitmap images. Nat. Methods. 11, 982–984 (2014).

52. K. I. Hng, D. Dormann, ConfocalCheck--a software tool for the automated monitoring of confocal microscope performance. PLoS One. 8, e79879 (2013).

53. E. Hill, Announcing the JCB DataViewer, a browser-based application for viewing original image files. J. Cell Biol. 183, 969– 970 (2008).

54. J. R. Swedlow, I. G. Goldberg, E. Brauner, P. K. Sorger, Informatics and Quantitative Analysis in Biological Imaging. Science. 300, 100–102 (2003).

55. E. Williams, J. Moore, S. W. Li, G. Rustici, A. Tarkowska, A. Chessel, S. Leo, B. Antal, R. K. Ferguson, U. Sarkans, A. Brazma, R. E. Carazo Salas, J. R. Swedlow, Image Data Resource: a bioimage data integration and publication platform. Nat. Methods. 14, 775 (2017).

56. S. Ram, J. Liu, A Semiotics Framework for Analyzing Data Provenance Research. Journal of Computing Science and Engineering. 2, 221–248 (2008).

57. S. Ram, J. Liu, A Semantic Foundation for Provenance Management. J. Data Semant. 1, 11–17 (2012).

58. Movincell Consortium, Multi-dimensional marine organism dataview. Movincell (2015), (available at http://movincell.org/).

59. A. Iudin, P. K. Korir, J. Salavert-Torres, G. J. Kleywegt, A. Patwardhan, EMPIAR: a public archive for raw electron microscopy image data. Nat. Methods. 13, 387–388 (2016).

60. J. Ellenberg, J. R. Swedlow, M. Barlow, C. E. Cook, U. Sarkans, A. Patwardhan, A. Brazma, E. Birney, A call for public archives for biological image data. Nat. Methods. 15, 849–854 (2018).

61. Allen Institute of Cell Science, Allen Cell Explorer. www.allencell.org/ (2018), (xavailable at https://www.allencell.org/).

62. T. Guilbert, Cochin Image Database. Cochin Image Database (2019), (available at https://imagerie.cochin.inserm.fr/sis4web/login.php).

63. M. Ellisman, S. Peltier, D. Orloff, W. Willy Wong, S. Penticoff, Center for Research in Biological Systems, Cell Image Library. www.cellimagelibrary.org (2019), (xavailable at http://www.cellimagelibrary.org/home).

64. Y. Tohsato, K. H. L. Ho, K. Kyoda, S. Onami, SSBD: a database of quantitative data of spatiotemporal dynamics of biological phenomena. Bioinformatics. 32, 3471–3479 (2016).

65. D. N. Orloff, J. H. Iwasa, M. E. Martone, M. H. Ellisman, C. M. Kane, The cell: an image library-CCDB: a curated repository of microscopy data. Nucleic Acids Res. 41, D1241–50 (2013).

66. U. Sarkans, W. Chiu, L. Collinson, M. C. Darrow, J. Ellenberg, D. Grunwald, J.-K. Hériché, A. Iuding, G. G. Martins, T. Meehan, K. Narayan, A. Patwardhan, M. Robert, G. Russell, H. R. Saibil, C. Strambio-De-Castillia, J. R. Swedlow, C. Tischer, V. Uhlmann, P. Verkade, M. Barlow, O. Bayraktar, E. Birney, C. Catavitello, C. Cawthorne, S. Wagner-Conrad, E. Duke, P. Paul-Gilloteaux, E. Gustin, M. Harkiolaki, P. Kankaanpää, T. Lemberger, J. McEntyre, J. Moore, A. W. Nicholls, S. Onami, H. Parkinson, M. Parsons, M. Romanchikova, N. Sofroniew, J. Swoger, N. Utz, L. Voortman, F. Wong, P. Zhang, G. J. Kleywegt, A. Brazma, REMBI: Recommended Metadata for Biological Images - realizing the full potential of the bioimaging revolution by enabling data reuse. Nat. Methods. ePub (2021),, doi:10.1038/s41592-021-01166-8.

67. A. Rigano, C. Strambio-De-Castillia, Minimum Information About Particle Tracking Experiments. Biosharing.org (2016), (available at https://biosharing.org/bsg-000671).

68. P. Masuzzo, MIACME 0.1. Cell Migration Standardization Organization (2017), (available at http://cellmigstandorg.github.io/MIACME/v0.1/spec/).

69. A. Rigano, U. Boehm, J. J. Chambers, N. Gaudreault, J. A. Pimentel, D. Sudar, P. Bajcsy, C. M. Brown, A. D. Corbett, O. Faklaris, J. Lacoste, A. Laude, G. Nelson, R. Nitschke, A. J. North, D. Grunwald, C. Strambio-De-Castillia, 4DN-BINA-OME (NBO) Tiered Microscopy Metadata Specifications - v2.00 - XLS Spreadsheet and Entity Relationship schemas (https://github.com/WU-BIMAC/NBOMicroscopyMetadataSpecs, 2021; https://zenodo.org/record/4682459).

70. P. P.-S. Chen, The entity-relationship model—toward a unified view of data. ACM Transactions on Database Systems (TODS). 1, 9–36 (1976).

71. H. Peng, Bioimage informatics: a new area of engineering biology. Bioinformatics. 24, 1827–1836 (2008).

72. C. Allan, J.-M. Burel, J. Moore, C. Blackburn, M. Linkert, S. Loynton, D. MacDonald, W. J. Moore, C. Neves, A. Patterson, M. Porter, A. Tarkowska, B. Loranger, J. Avondo, I. Lagerstedt, L. Lianas, S. Leo, K. Hands, R. T. Hay, A. Patwardhan, C. Best, G. J. Kleywegt, G. Zanetti, J. R. Swedlow, OMERO: flexible, model-driven data management for experimental biology. Nat. Methods. 9, 245–253 (2012).

73. A. Rigano, U. Boehm, J. J. Chambers, N. Gaudreault, A. J. North, J. A. Pimentel, D. Sudar, P. Bajcsy, C. M. Brown, A. D. Corbett, O. Faklaris, J. Lacoste, A. Laude, G. Nelson, R. Nitschke, D. Grunwald, C. Strambio-De-Castillia, 4DN-BINA-OME (NBO) Tiered Microscopy Metadata Specifications - v2.01 - XLS Spreadsheet and Entity Relationship schemas (https://github.com/WU-BIMAC/NBOMicroscopyMetadataSpecs, 2021; https://zenodo.org/record/4711229).

74. K. Hoffmann, U. Resch-Genger, R. Nitschke, in Standardization and Quality Assurance in Fluorescence Measurements II (Springer, 2008; https://link.springer.com/chapter/10.1007/4243_2008_028), pp. 89–116.

75. U. Resch-Genger, K. Hoffmann, W. Nietfeld, A. Engel, J. Neukammer, R. Nitschke, B. Ebert, R. Macdonald, How to improve quality assurance in fluorometry: fluorescence-inherent sources of error and suited fluorescence standards. J. Fluoresc. 15, 337– 362 (2005).

76. G. Nelson, L. Gelman, O. Faklaris, R. Nitschke, A. Laude, Interpretation of Confocal ISO 21073: 2019 confocal microscopes: Optical data of fluorescence confocal microscopes for biological imaging-Recommended Methodology for Quality Control. arXiv [q-bio.OT] (2020), (available at http://arxiv.org/abs/2011.08713).

77. R. P. J. Nieuwenhuizen, K. A. Lidke, M. Bates, D. L. Puig, D. Grunwald, S. Stallinga, B. Rieger, Measuring image resolution in optical nanoscopy. Nat. Methods. 10, 557–562 (2013).

78. R. W. Cole, T. Jinadasa, C. M. Brown, Measuring and interpreting point spread functions to determine confocal microscope resolution and ensure quality control. Nat. Protoc. 6, 1929–1941 (2011).

79. R. W. Cole, M. Thibault, C. J. Bayles, B. Eason, A.-M. Girard, T. Jinadasa, C. Opansky, K. Schulz, C. M. Brown, International test results for objective lens quality, resolution, spectral accuracy and spectral separation for confocal laser scanning microscopes. Microsc. Microanal. 19, 1653–1668 (2013).

80. B. Eason, D. Young, A. J. Spurmanis, T. L. E. Wee, D. Kaufman, C. M. Brown, in Microscopy: advances in scientific research and education, A. Mendez-Vilas, Ed. (Formatex Research Center, 2014), pp. 713–724.

81. M. Halter, E. Bier, P. C. DeRose, G. A. Cooksey, S. J. Choquette, A. L. Plant, J. T. Elliott, An automated protocol for performance benchmarking a widefield fluorescence microscope. Cytometry A. 85, 978–985 (2014).

82. ASTM International, “ASTM F3294 - 18 Standard Guide for Performing Quantitative Fluorescence Intensity Measurements in Cell-based Assays with Widefield Epifluorescence Microscopy” (ASTM F3294, 2018),, doi:https://doi.org/10.1520/F3294-18.

83. M. A. Model, J. K. Burkhardt, A standard for calibration and shading correction of a fluorescence microscope. Cytometry. 44, 309–316 (2001).

84. R. M. Zucker, O. T. Price, Statistical evaluation of confocal microscopy images. Cytometry. 44, 295–308 (2001).

85. R. M. Zucker, O. Price, Evaluation of confocal microscopy system performance. Cytometry. 44, 273–294 (2001).

86. E. H. Cho, S. J. Lockett, Calibration and standardization of the emission light path of confocal microscopes. J. Microsc. 223, 15– 25 (2006).

87. S. Bradner, “[RFC2119] Key words for use in RFCs to Indicate Requirement Levels” (Internet RFC 2119, 1997), (available at http://www.ietf.org/rfc/rfc2119.txt).

88. J. M. Heddleston, J. S. Aaron, S. Khuon, T.-L. Chew, A guide to accurate reporting in digital image processing: can anyone reproduce your quantitative analysis? J. Cell Sci. 134: jcs254151 (2021), doi:10.1242/jcs.254151.

89. J. S. Aaron, T.-L. Chew, A guide to accurate reporting in digital image acquisition: can anyone replicate your microscopy data? J. Cell Sci. 134: jcs254144 (2021), doi:10.1242/jcs.254144.

90. P. Montero Llopis, R. A. Senft, T. J. Ross-Elliott, R. Stephansky, D. P. Keeley, P. Kosha, G. Marqués, Y. Gao, B. R. Carlson, T. Pengo, M. A. Sanders, L. A. Cameron, M. S. Itano, Best practices and tools for reporting reproducible fluorescence microscopy methods. Nat. Methods. 35 (2021), doi:10.1038/s41592-021-01156-w.

91. A. Rigano, A. Balashov, S. Ehmsen, S. U. Ozturk, B. Alver, C. Strambio-De-Castillia, Micro-Meta App - Electron (Github - https://github.com/WU-BIMAC, 2021; https://doi.org/10.5281/zenodo.4750765).

92. J. Rung, A. Brazma, Reuse of public genome-wide gene expression data. Nat. Rev. Genet. 14, 89–99 (2013).

93. A. Brazma, P. Hingamp, J. Quackenbush, G. Sherlock, P. Spellman, C. Stoeckert, J. Aach, W. Ansorge, C. A. Ball, H. C. Causton, T. Gaasterland, P. Glenisson, F. C. Holstege, I. F. Kim, V. Markowitz, J. C. Matese, H. Parkinson, A. Robinson, U. Sarkans, S. Schulze-Kremer, J. Stewart, R. Taylor, J. Vilo, M. Vingron, Minimum information about a microarray experiment (MIAME)-toward standards for microarray data. Nat. Genet. 29, 365–371 (2001).

94. J. P. A. Ioannidis, D. B. Allison, C. A. Ball, I. Coulibaly, X. Cui, A. C. Culhane, M. Falchi, C. Furlanello, L. Game, G. Jurman, J. Mangion, T. Mehta, M. Nitzberg, G. P. Page, E. Petretto, V. van Noort, Repeatability of published microarray gene expression analyses. Nat. Genet. 41, 149–155 (2009).

95. A. Brazma, Minimum Information About a Microarray Experiment (MIAME)--successes, failures, challenges. ScientificWorldJournal. 9, 420–423 (2009).

96. S.-A. Sansone, P. Rocca-Serra, M. Brandizi, A. Brazma, D. Field, J. Fostel, A. G. Garrow, J. Gilbert, F. Goodsaid, N. Hardy, P. Jones, A. Lister, M. Miller, N. Morrison, T. Rayner, N. Sklyar, C. Taylor, W. Tong, G. Warner, S. Wiemann, The First RSBI (ISA-TAB) Workshop: “Can a Simple Format Work for Complex Studies?” OMICS. 12, 143–149 (2008).

97. EMBL-EBI BioImage Archive. EMBL-EBI BioImage Archive (2019), (available at https://www.ebi.ac.uk/bioimage-archive/).

98. ISO, BBMRI-ERIC, “ISO/WD TS 23494-1 - Biotechnology — Provenance information model for biological material and data — Part 1: Design concepts and general requirements” (ISO/WD TS 23494–1, 2020), (available at https://www.iso.org/standard/80715.html).

99. L. Allen, A. O’Connell, V. Kiermer, How can we ensure visibility and diversity in research contributions? How the Contributor Role Taxonomy (CRediT) is helping the shift from authorship to contributorship. Learn. Publ. 32, 71–74 (2019).

100. Elsevier editors, CRediT author statement - editorial. Elsevier (2020) (available at https://www.elsevier.com/authors/policies-and-guidelines/credit-author-statement).

101. W3 Consortium, RDF Schema 1.1. w3.org (2014), (available at https://www.w3.org/TR/rdf-schema/).

102. G. Klyne, J. J. Carroll, Resource description framework (RDF): Concepts and abstract syntax. w3.ort (2014), (available at https://www.w3.org/TR/rdf-concepts/).

103. D. L. McGuinness, F. Van Harmelen, Others, OWL web ontology language overview. w3.org (2004), (available at https://www.w3.org/TR/owl-features/).

104. N. Kobayashi, S. Kume, K. Lenz, H. Masuya, RIKEN MetaDatabase: A Database Platform for Health Care and Life Sciences as a Microcosm of Linked Open Data Cloud. IJSWIS. 14, 140–164 (2018).

105. S. Kume, H. Masuya, M. Maeda, M. Suga, Y. Kataoka, N. Kobayashi, in Semantic Technology (Springer International Publishing, 2017; http://dx.doi.org/10.1007/978-3-319-70682-5_19), pp. 277–285.

106. S. Kume, H. Masuya, Y. Kataoka, N. Kobayashi, in Proceedings of the15th International Semantic Web Conference (Posters & Demos), P. Groth, E. Simperl, A. Gray, M. Sabou, M. Krötzsch, F. Lecue, F. Flöck, Y. Gil, Eds. (researchgate.net, 2016; http://ceur-ws.org/Vol-1690/paper93.pdf).

107. M. Hammer, N. Kobayashi, J. Moore, A. Rigano, F. Farzam, S. Onami, J. R. Swedlow, D. Grünwald, C. Strambio-De-Castillia, Metadata and Performance Tracking for Fluorescent Microscopes - Towards a Shared Microscopy Ontology - The 4DN-OME ontology: an OME-OWL extension with emphasis on usability, minimum information guidelines and quality control for super-resolution fluorescence microscopy (December, 5-6 2019),, doi:10.6084/m9.figshare.12299915.

