## Supplemental Material for "Towards community-driven metadata standards for light microscopy: tiered specifications extending the OME model"

-------------------------

#### SUPPLEMENTAL MATERIAL

#### Online resources

1. A. Rigano, U. Boehm, J.J. Chambers, N. Gaudreault, A.J. North, J.A. Pimentel, D. Sudar, P. Bajcsy, C.M. Brown, A.D. Corbett, O. Faklaris, J. Lacoste, A. Laude, G. Nelson, R. Nitschke, D. Grunwald, C. Strambio-De-Castillia (2021). **NBOMicroscopyMetadataSpecs: 4DN-BINA-OME (NBO) Microscopy Metadata Specifications.** *Available at:* *https://zenodo.org/record/4710731*

2. A. Rigano, U. Boehm, J.J. Chambers, N. Gaudreault, A.J. North, J.A. Pimentel, D. Sudar, P. Bajcsy, C.M. Brown, A.D. Corbett, O. Faklaris, J. Lacoste, A. Laude, G. Nelson, R. Nitschke, D. Grunwald, C. Strambio-De-Castillia (2021). **4DN-BINA-OME (NBO) Tiered Microscopy Metadata Specifications - v2.01 - XLS Spreadsheet and Entity Relationship schemas.** *Available at: https://zenodo.org/record/4711229*

3. A. Rigano and C. Strambio-De-Castillia (2021). **4DN-BINA-OME (NBO) Tiered Microscopy Metadata Specifications - v2.01 - XSD schema.** *Available at: https://doi.org/10.5281/zenodo.4711426*

#### Supplemental Figures

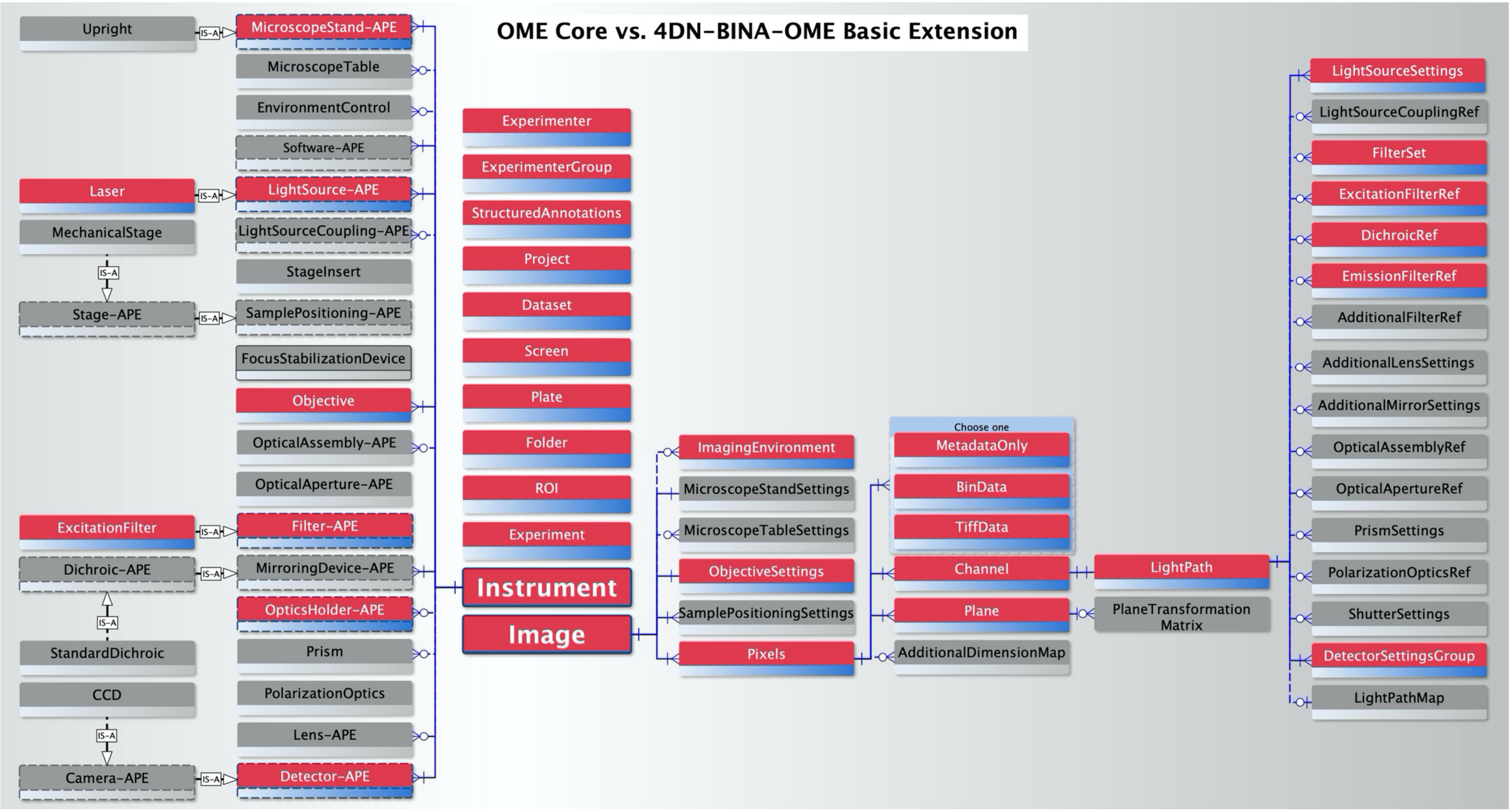


**Supplemental Figure 1: A) Comparison between the Core of the OME Data Model and the 4DN-BINA Basic Extension.** Simplified Entity-Relationship diagrammatic representation of the Basic extension of the ***<INSTRUMENT>*** and ***<IMAGE>*** elements of the OME Data Model that is put forth by the 4D Nucleome (4DN) Imaging Standards Working Group (IWG) (1,2) and by the Bioimaging North America (BINA) Quality Control and Data Management Working Group (QC-DM-WG) (3). *Red and Blue* boxes represent elements that belong to the Core of the OME Data Model. *Grey* boxes represent elements that were introduced *de novo* as part of the proposed extension. Dashed-lined boxes represent *Abstract Parent Elements (APE*). These elements describe general families to which the indicated "children" elements belong. For example, in the model a ***<Laser>*** *IS-A* (*black dashed arrow*) child of a ***<LightSource-APE>***. For simplicity's sake only a selected subset of all available children-elements is indicated in this summary diagram.

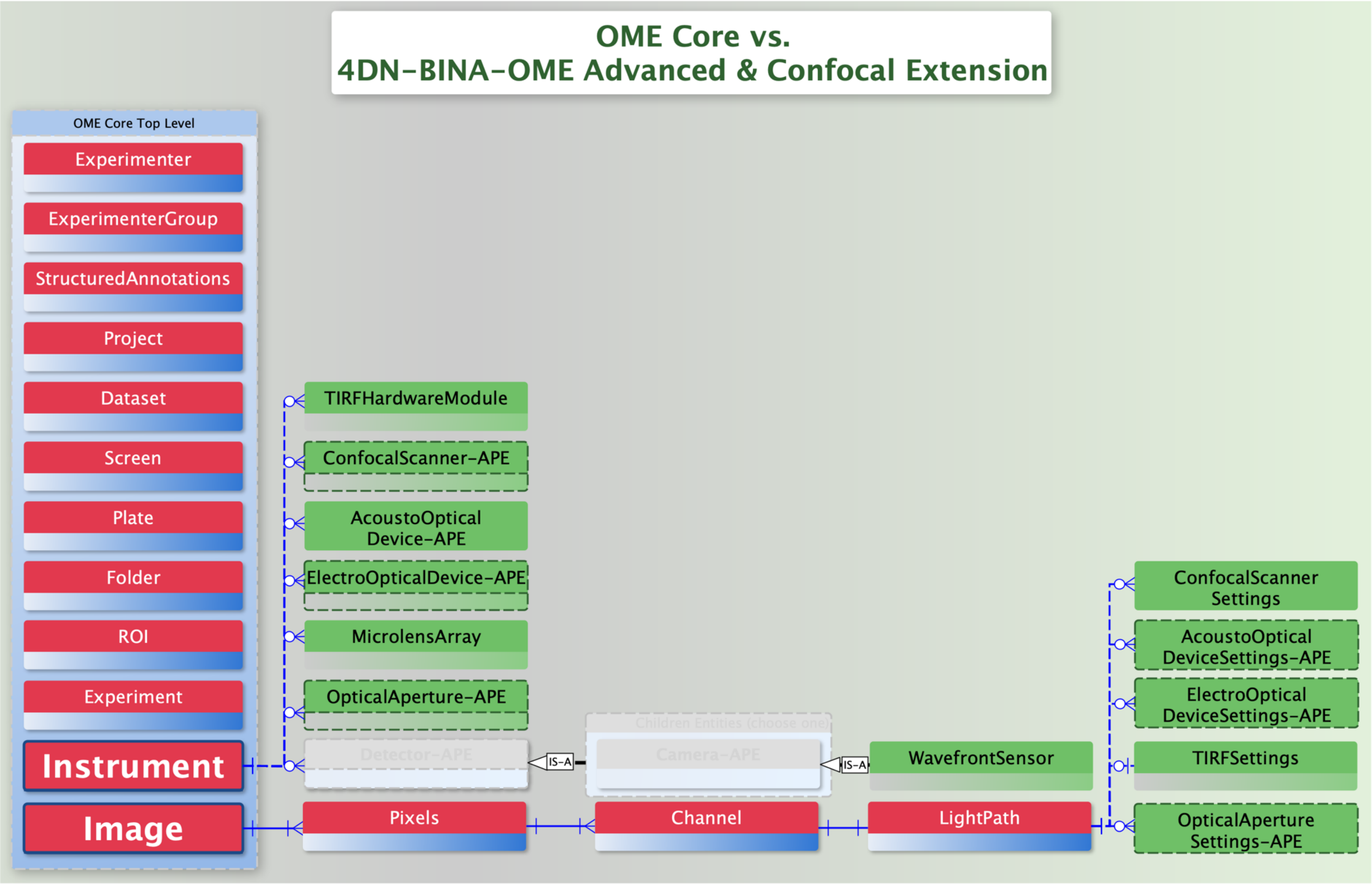


**Supplemental Figure 1: B) Comparison between the Core of the OME Data Model and the 4DN-BINA Advanced & Confocal Extension.** Simplified Entity-Relationship diagrammatic representation of the Advanced & Confocal extension of the ***<INSTRUMENT>*** and ***<IMAGE>*** elements of the OME Data Model that is put forth by 4DN and BINA (1–3). *Red and Blue* boxes represent elements that belong to the Core of the OME Data Model. *Green* boxes represent elements that were introduced *de novo* as part of the proposed extension. Dashed-lined boxes represent *Abstract Parent Elements (APE)*. These elements describe general families to which the indicated "children" elements belong. For example, in the model a **<WaveFrontSensor>** *IS-A* (*black dashed arrow*) child of a ***<Camera-APE>***. For simplicity's sake only a selected subset of all available children-elements is indicated in this summary diagram.

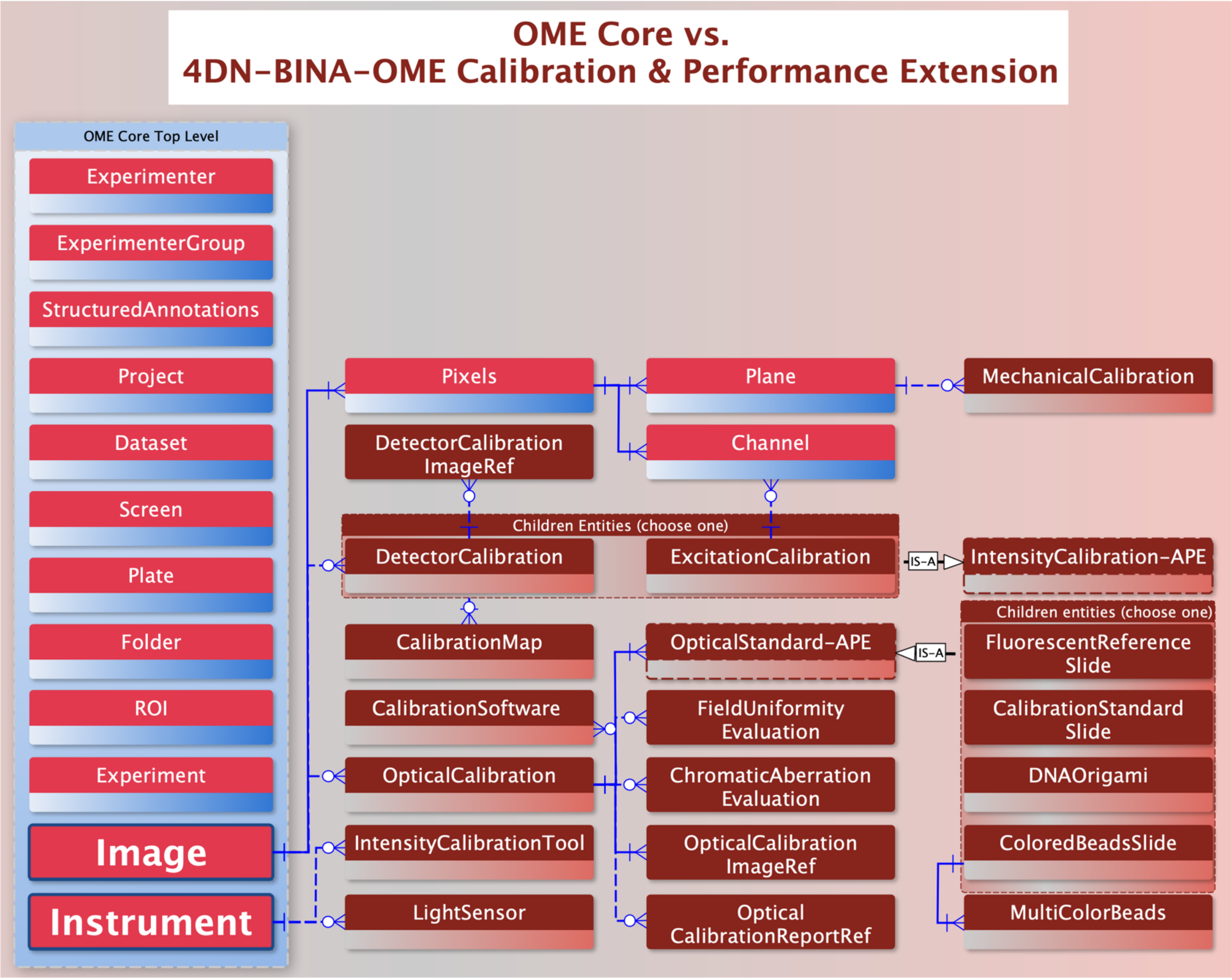


**Supplemental Figure 1:C) Comparison between the Core of the OME Data Model and the 4DN-BINA Calibration & Performance Extension.** Simplified Entity-Relationship diagrammatic representation of the Calibration & Performance extension of the ***<INSTRUMENT>*** and ***<IMAGE>*** elements of the OME Data Model that is put forth by 4DN and BINA (1–3). *Red and Blue* boxes represent elements that belong to the Core of the OME Data Model. *Maroon* boxes represent elements that were introduced *de novo* as part of the proposed extension. Dashed-lined boxes represent *Abstract Parent Elements (APE)*. These elements describe general families to which the indicated "children" elements belong. For example, in the model a ***<CalibrationStandardSlide>*** *IS-A* (*black dashed arrow*) child of a ***<OpticalStandard-APE>***. For simplicity's sake only a selected subset of all available children-elements is indicated in this summary diagram.

### OME Core vs. 4DN-BINA-OME Basic Extension Microscope HARDWARE Specifications

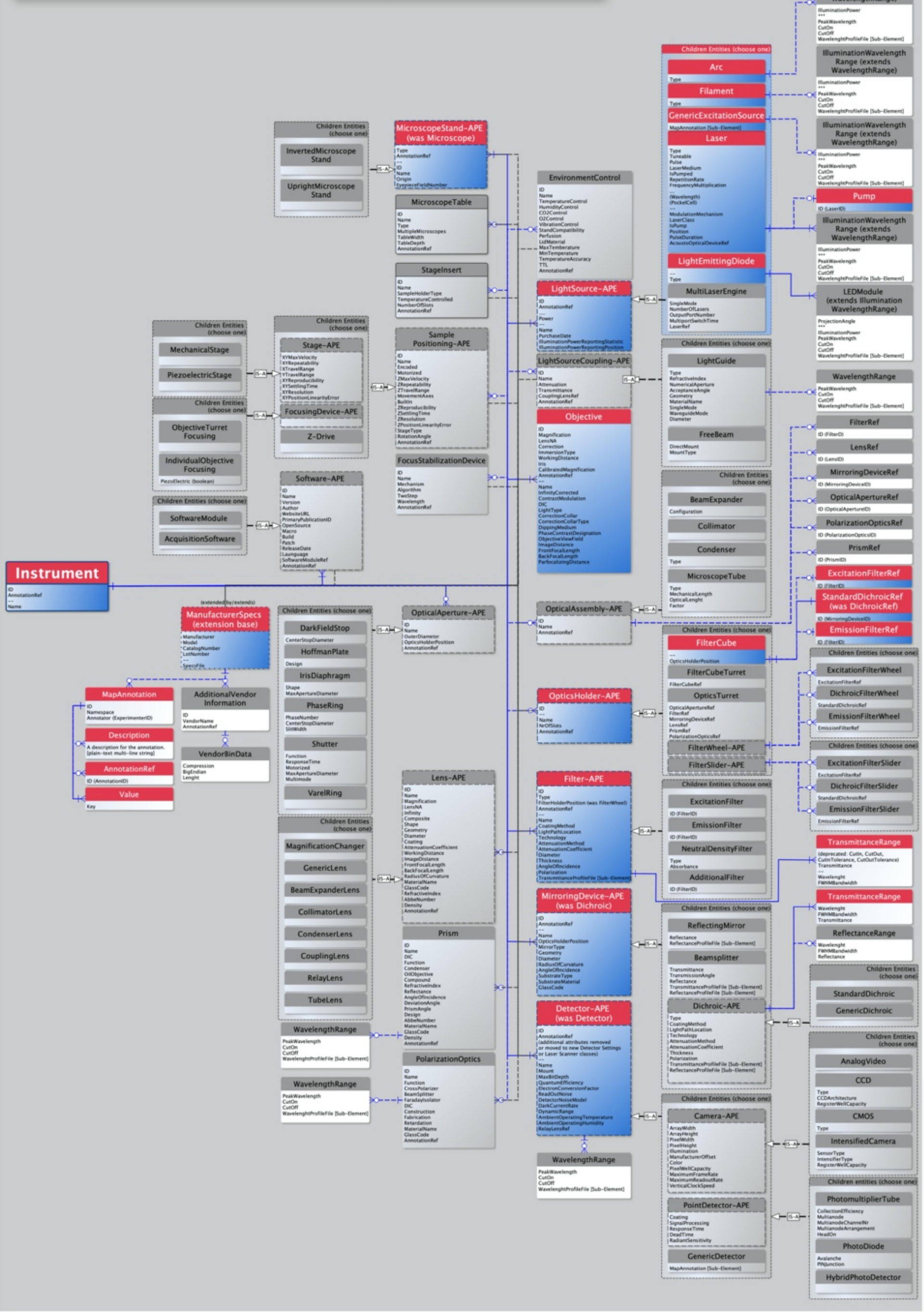


### OME Core vs. 4DN-BINA-OME Basic Extension Image ACQUISITION Settings

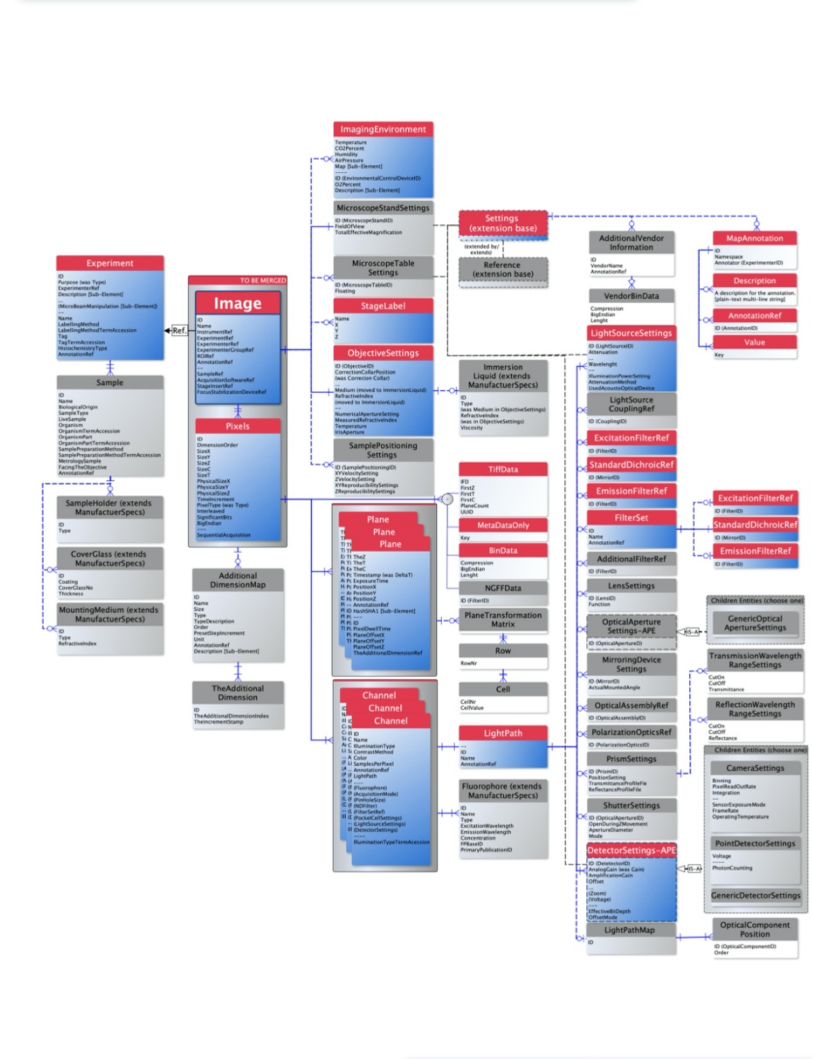


#### Legend

This is a Entity-Relationship diagrammatic representation of a proposed REVISION of the OME Core data model (blue/red boxes) along side a proposed OME BASIC EXTENSION (grey boxes) developed by members of the Imaging Working Group of the 4D Nucleome network (https://www.4dnucleome.org) and by members of the BINA Quality Control and Data Management Working Group (https://www.bioimagingna.org/qc-dm-wg).

The Entity-Relationship formalism can be utilized to model information about a real-world situation or an actual object (in this case a Microscope INSTRUMENT and an IMAGE acquired using that Instrument) by using three types of model elements:
1) ENTITIES, represented by Boxes;
2) RELATIONSHIPS, represented by blue lines connecting boxes; and
3) ATTRIBUTES, represented by fields within boxes

When describing a real-life situation or object:
1) ENTITIES correspond to NOUNS = the items we want to collect information about.
2) RELATIONSHIPS correspond to VERBS = actions/states/occurrences that connect Entities with each other
3) ATTRIBUTES correspond to ADJECTIVES = they describe the actual information (in our case metadata) that we want to record about each Entity.

In order to interpret the schema please start either from the <INSTRUMENT> element for the Microscope Hardware Specifications section of the diagram or from the <IMAGE> element of the Image Acquisition Settings sections.Then follow the blue lines to the connected boxes and think something like:
1) An Instrument has a Microscope Body, might rest on a Microscope Table, utilizes a Light Source and magnifies the sample using an Objective.
2) Alternatively, an Image was produced as part of a specific Experiment, was collected in a specific Imaging Environment and using specific Microscope Settings and has four Channels.

For questions or comments please contact:

#### Notes

Blue edges, blue edges represent RELATIONSHIPs between entities. SOLID EDGES are used to connect an Element with a REQUIRED Sub-Element. DASHED EDGED are used to connect an Element with an OPTIONAL Sub-Element.
In addition, different CARDINALITIES can be represented as follow:

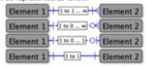


Black dashed arrows, these arrows simbolize an inheritance relationship similar to what would connect a hypothetical "Feline" parent element with a "Cat" child element. In this example, while the "Cat" element represents a "concrete" entity that exist in the real-world, the "Feline" element represents an "abstract" category that is only useful for categorization pruposes. In a similar manner, these arrows connect concrete elements that represents actual microscope hardware components with their Abstract Parent Elements (APE), which are utilized for making the model less repetitive and more efficient to read by software tools.
These arrows should be read as follows, a "Laser" IS-A "LightSource".

Dashed grey edges, these edges signify "Extends" and they should be read as follows: "Objective" extends "ManufacturerSpecs". In this example, the concept of extension indicates that Objective is of type "ManufacturerSpecs" but it has additional attributes that are unique to it and distinguishes from other elements.
-
-AnnotationRef, This element always refers to a Comment/Annotation element as described for Channel. However for simplicity sake most Comment/Annotation elements have been omitted and the AnnotationRef has been inserted in the referring element as an attribute.
-[Sub-Element], For simplicity and when indicated Sub-Elements are listed within the referring element as an attribute.
-References, For simplicity and when indicated, Reference elements (e.g., ImageRef) are listed whithin the referring element as attributes.
-Units were omitted from this diagram for simplicity. However, they are essential for reproducibility and they are included in the model and in all its formal representations.
-Attributes listed after the --- separator have been added to the OME Core as part of the proposed revision.
-Attributes listed in parenthesis and after the .... separator have been removed as part of the proposed extension.
-APE, Abstract Parent Entity (see explanation above).

**Supplemental Figure 2: Entity-Relationship diagram representing the BASIC 4DN-BINA extension of the OME Data Model**. This is an Entity-Relationship diagrammatic representation of the BASIC EXTENSION (grey boxes) of the OME Core Data Model (blue/red boxes) proposed by 4DN and BINA (1–3). Downloadable version of this schema is available online (7,69). The Entity-Relationship formalism can be utilized to model information about a real-world situation or actual objects (in this case a microscope INSTRUMENT and an IMAGE acquired using that Instrument) by using three types of model elements: 1) Entities, represented by Boxes; 2) Relationships, represented by blue lines connecting boxes; and 3) Attributes, represented by fields within boxes. When describing a real-life situation or an object, entities can be thought of as nouns that represent the items that we want to collect information about. At the same time relationships are analogous to verbs and represent actions/states/occurrences that connect Entities with each other. Finally, attributes correspond to adjectives and they describe the actual information (in our case metadata) about each Entity we want to collect. To interpret the schema please start from either the ***<INSTRUMENT>*** or **<IMAGE>** elements of the Microscope Hardware Specifications and Image Acquisition Settings sections of the diagram, respectively. Then follow the blue lines to the connected boxes and think something like, an Instrument has a Microscope Body, might rest on a Microscope Table, utilizes a Light Source, and magnifies the sample using an Objective. Alternatively, an Image was produced as part of a specific Experiment, was collected in a specific Imaging Environment using specific Microscope Settings and has four Channels. Black dashed arrows symbolize an inheritance relationship like what would connect a hypothetical "Feline" parent element with a "Cat" child element. In this example, while the "Cat" element represents a "concrete" entity that exists in the real world, the "Feline" element represents an "abstract" category that is only useful for categorization purposes. In a similar manner, these arrows connect concrete elements that represent actual microscope hardware components, with their Abstract Parent Elements (APE), which are useful for making the model less repetitive and more efficient to read by software tools. These arrows should be read as follows, a "Laser" IS-A "LightSource". Grey dashed edges signify "Extends" and they should be read as follows: "Objective" extends "ManufacturerSpecs". Units were omitted from this diagram for simplicity. However, they are essential for reproducibility and they are included in the model and in all its formal representations. Attributes listed after the --- separator have been added to the OME Core as part of the proposed revision. Attributes listed in parenthesis and after the .... separator have been removed as part of the proposed extension. Abbreviations: APE, Abstract Parent Entity (see explanation above).

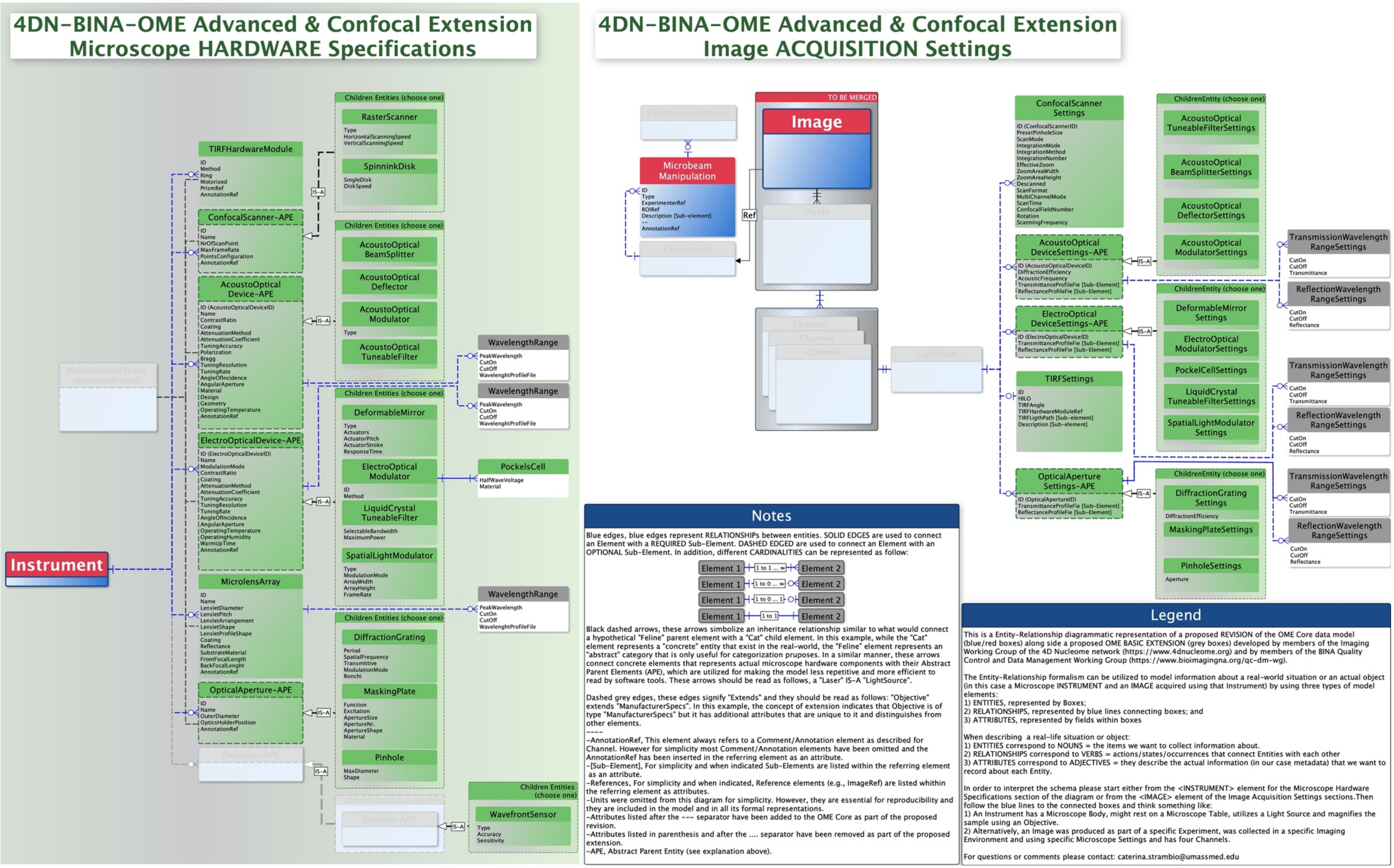


**Supplemental Figure 3: Entity-Relationship diagram representing the ADVANCED & CONFOCAL 4DN-BINA extension of the OME Data Model**. This is an Entity-Relationship diagrammatic representation of the ADVANCED & CONFOCAL EXTENSION (grey boxes) of the OME Core Data Model (blue/red boxes) proposed by 4DN and BINA (1–3). Downloadable version of this schema is available online (7,69). For more details, please see the legend for Supplemental Figure 1.

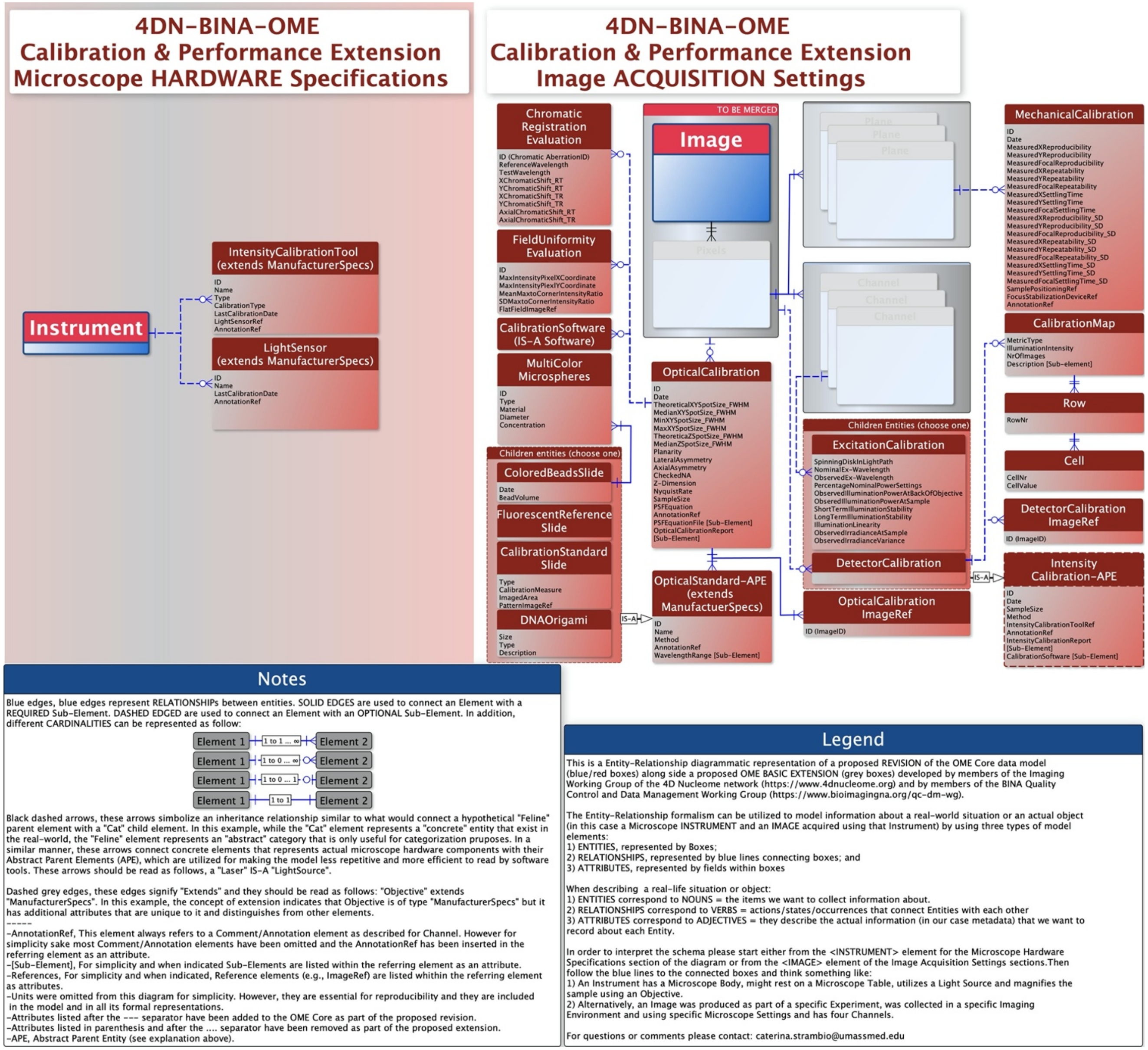


**Supplemental Figure 4: Entity-Relationship diagram representing the CALIBRATION & PERFORMANCE 4DN-BINA extension of the OME Data Model**. This is an Entity-Relationship diagrammatic representation of the CALIBRATION & PERFORMANCE EXTENSION (maroon boxes) of the OME Core Data Model (blue/red boxes) proposed by 4DN and BINA (1–3). Downloadable version of this schema is available online (7,69). For more details, please see the legend for Supplemental Figure 1.

### Supplemental Tables

### Supplemental Table I

**Supplemental Table I: Extended view of the Tiers for the reporting of Light Microscopy Metadata as proposed by the Imaging Standards Working Group of the 4D Nucleome initiative and by the Quality Control and Data Management Working Group of Bioimaging North America.**

| Category | Tier Nr. | Name | Description | Example Experiment Type | Example Labelling | Experimental/ Sample | Microscope hardware specifications | Image acquisition settings | Optical calibration | Intensity calibration | Mechanical calibration |
|---|---|---|---|---|---|---|---|---|---|---|---|
| Descriptive | 1 | Minimum Information/ Material & Methods | Reporting qualitative effects, or effects that require simple quantification including the identification of non-refractive limited objects followed by basic feature extraction and statistical analysis | Transfection control, viability assay, counting of cells and nuclei, expression level measurements, localization of markers in cellular sub-compartments | Fluorescent In Situ Hybridization (FISH), Immuno Fluorescence (IF), Fluoresent Protein (FP) labelling | experimenter name; experiment description and date; sample description; mounting medium; temperature and CO2 conditions; | microscope manufacturer, model and type; light source manufacturer, wavelenght and type; objective manufacturer, magnification, NA and correction; filter/dichroic transmittance range; detector manufacturer and type | acquisition date; immersion liquid name and refractive index; illumination type and intensity; fluorophore; exposure time; pixel dwell time; channel name, color, contrast method and acquisition mode; image dimension order and number; physical pixel size x, y, and z | not required; recommended quarterly | not required; recommended annually | |
| Analytical | 2 | Advanced Quantification and/or Live Cell Imaging | Identification and localization of diffraction-limited particles, super-resolution microscopy, tracking of intracellular dynamics | Diffraction-limited spot localization, measurement of distances, co-localization studies, signal-starved features, advanced processing, cell tracking and single-particle tracking, dynamic expression level quantification | All of the above + Single Molecule (SM) FISH, CasFISH, SM Proximity Ligation Assay (PLA), dCas9-based labelling, OligoPaint | O2 pressure, and humidity conditions; refractive index of the mounting medium; thickness of the coverglass | detailed environmental control device, microscope table, light source, light source coupling, transmittance light path, magnification, sample positioning, focusing, autofocus, filter, dichroic, additional optics and detector specification (e.g., lightsource spectral properties; objective correction properties; focusing device ZReproducibility, ZSettlingTime, ZResolution, etc.) | illumination attenuation; objective temperature and iris aperture; immersion liquid measured refractive index; sample positioning settings; detector integration; ligthpath configuration | required monthly | highly-recommended monthly to quarterly | |
| | 3 | Manufacturing/ Technical Development/ Full Documentation | Full documentation of microscopic setup, image acquisition and quality control | Microscopy hardware manufacturing; development of novel unproven technology in both commercial and academic settings; full reproducibility of microscopy set-up and image acquisition settings | All of the above | all the metadata specified by the data model - including any novel technology-specific metrics | | | required for every acquisition | required monthly to quarterly | |

**Legend:** Each tier accommodates increasingly complex images, experiments, instrumentation, and analytical needs and therefore requires progressively more metadata. This tiered system is not intended to meet the needs of all imaging communities. Rather it is proposed as a framework that might need to be adapted and modified depending on the needs of individual data collection consortia, disciplines, or institutions.
